# EDoF-Miniscope: pupil engineering for extended depth-of-field imaging in a fluorescence miniscope

**DOI:** 10.1101/2022.08.05.502947

**Authors:** Joseph Greene, Yujia Xue, Jeffrey Alido, Alex Matlock, Guorong Hu, Kivilcim Kiliç, Ian Davison, Lei Tian

## Abstract

Extended depth of field (EDoF) microscopy has emerged as a powerful solution to greatly increase the access into neuronal populations in table-top imaging platforms. Here, we present EDoF-Miniscope, which integrates an optimized thin and lightweight binary diffractive optical element (DOE) onto the gradient refractive index (GRIN) lens of a head-mounted fluorescence miniature microscope, i.e. “miniscope”. We achieve an alignment accuracy of 70 μm to allow a 2.8X depth-of-field extension between the twin foci. We optimize the phase profile across the whole back aperture through a genetic algorithm that considers the primary GRIN lens aberrations, optical property of the submersion media, and axial intensity loss from tissue scattering in a Fourier optics forward model. Compared to other computational miniscopes, our EDoF-Miniscope produces high-contrast signals that can be recovered by a simple algorithm and can successfully capture volumetrically distributed neuronal signals without significantly compromising the speed, signal-to-noise, signal-to-background, and maintain a comparable 0.9-μm lateral spatial resolution and the size and weight of the miniature platform. We demonstrate the robustness of EDoF-Miniscope against scattering by characterizing its performance in 5-μm and 10-μm beads embedded in scattering phantoms. We demonstrate that EDoF-Miniscope facilitates deeper interrogations of neuronal populations in a 100-μm thick mouse brain sample, as well as vessels in a mouse brain. Built from off-the-shelf components augmented by a customizable DOE, we expect that this low-cost EDoF-Miniscope may find utility in a wide range of neural recording applications.

## Introduction

Head-mounted miniaturized fluorescence microscopes, or miniscopes, have become an invaluable tool for interrogating neural activity in freely behaving animals by employing 3D printing and off-the-shelf miniature optics to enable low-cost one-photon (1P) neural imaging across various in-vivo studies^1^. Many existing miniscopes incorporate high numerical aperture (NA; e.g. 0.5-0.55) gradient refractive index (GRIN) objective lenses^2–4^ that limit the depth of field (DoF) to a shallow region near the end surface of the lens^1^. However, neurons are inherently distributed in 3D, which leads to a *need* of probing an extended depth range to facilitate studies on large neural populations. Miniscopes incorporating electrowetting lenses^5^ enable focus adjustment; however they have to trade frame rate for reaching deeper structures. This leads to an undesirable tradeoff between capturing high-speed neural dynamics and the number of accessible depths. Miniaturized lightfield miniscopes circumvent the shallow DoF by encoding 3D information using a microlens array^6–8^; however, they have to trade off the signal-to-noise ratio (SNR), signal-to-background ratio (SBR), and spatial resolution due to high-degree of optical multiplexing. In general, 1P miniscopes will benefit from an extended depth of field (EDoF) imaging technique that can record from a volumetrically distributed neural population without significantly compromising the speed, SNR, SBR, and spatial resolution.

In the past few decades, many computational microscopy systems have been developed to record fluorescence signals from an extended depth range. Depending on the final goal, one can classify these techniques into two main categories including “2D-to-3D” and “2D-to-2D” techniques. The “2D-to-3D” techniques explicitly reconstruct 3D fluorescence from a 2D image. Miniaturized computational single-shot 3D fluorescence microscopes, such as MiniLFM^6^, Miniscope3D^8^, Bio-FlatScope^9^, GEOMscope^10^, and Computational Miniature Mesoscope^11^, encode depth information using a microlens array, customized phase mask or amplitude mask, and require computationally intensive 2D-to-3D model-based^8–11^ or deep-learning^12,13^ deconvolution algorithms to recover the 3D fluorescence distribution.

Alternatively, the “2D-to-2D” EDoF techniques “compress” fluorescence signals from an extended depth into a 2D image while neglecting the accurate depth information, and has been shown to increase the access in neural imaging without costly inverse algorithms in several *tabletop* platforms^14–17^. The EDoF capability is typically achieved by pupil engineering, which uses a custom component to modify the optical field as it passes through the pupil plane of a microscope. A large family of pupil functions have been shown to achieve an EDoF^18–21^. However, many of these systems rely on active optical devices, such as spatial light modulators (SLM), to project the desired function onto the pupil plane of the objective lens. These devices exhibit large form factors and bulky adapters that limit their integration into miniaturized platforms^22^. Alternatively, diffractive optical elements (DOEs) have been used to directly shape the wavefront at the pupil plane as a compact, lightweight alternative for pupil engineering^23,24^.

Our goal here is to employ phase-only DOE-based pupil engineering to achieve high-contrast EDoF imaging and directly integrate into a miniscope. To this end, we develop EDoF-Miniscope (**Fig. 1A-C**) that extends the recoverable depth from ∼18 μm to ∼51 μm (**Fig. 1D**) while maintaining the spatial resolution (**Fig. 1F-G**). By employing 2D-to-2D encoding, EDoF-Miniscope only requires a simple wavelet filter^25^ for real-time post processing to facilitate neural signal extraction (**Fig. 1H-K**). For ease of fabrication, we employ a binary phase DOE that is fabricated by a single-step photolithography (**Fig. 1C**). We employ a genetic algorithm to optimize the binary phase mask from a basis of three widely used EDoF phase functions, consisting of axicon, spherical aberration, and defocus^26^ (**Fig. 1E**). As compared to gradient-based optimization strategies, such as those based on deep learning^23,27,28^, our genetic algorithm allows us to more flexibly incorporate non-differentiable constraints, such as constraining relative peak intensity over the desired EDoF range and binarization of the phase profile. Our algorithm is built on a linear shift invariant Fourier-optics model and, in addition, incorporates the native spherical aberration of the GRIN lens and the scattering-induced intensity decay. As a result, the optimized binary phase profile achieves an EDoF in addition to the native spherical aberration-induced point spread function (PSF) elongation and is robust to signal loss due to scattering. Furthermore, the defocus term axially displaces the PSF so that both the +1^st^ and -1^st^ order foci are within the focal region (**Fig. 1D**). In this way, our design further extends the EDoF range.

**Fig. 1.**
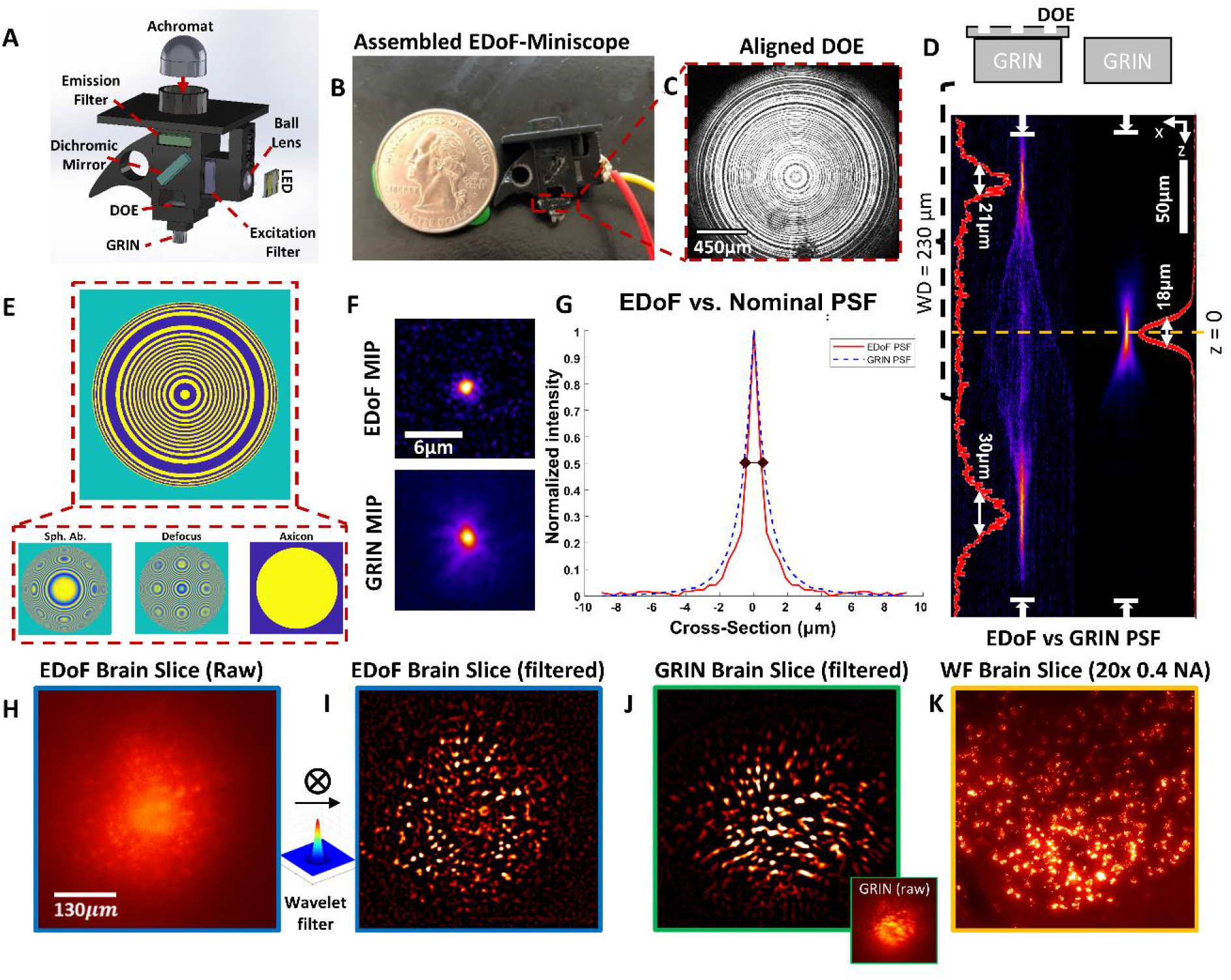
Pupil engineering for EDoF miniscope. (**A**) The EDoF-Miniscope combines a lightweight DOE into a miniscope. (**B**) Picture of the EDoF-Miniscope prototype (sensor omitted). (**C**) Image of the DOE aligned and glued to the back of the 0.55 NA GRIN lens with a collimated laser illumination. (**D**) XZ cross-section of the 3D PSFs for EDoF-Miniscope (left) and miniscope (right) captured by scanning a 1-μm fluorescent bead on a glass slide through a 346-μm depth range with a 1-μm step size. The GRIN lens working distance (WD; 0.23mm) allows utilizing both the 1^st^ (bottom) and -1^st^ (top) diffraction orders created by the DOE to extend the PSF. The Full Width Half Maximum (FWHM) of the GRIN PSF is 18 μm and the FWHM of the EDoF-Miniscope PSF is 30 μm (+1st order) and 21 μm (−1 order) yielding a 2.8x improvement in the DoF. (**E**) Breakdown of the optimized phase functions that comprise the DOE. (**F**) XY MIP of the EDoF-Miniscope (top) and miniscope (bottom) PSF over their focal range. (**G**) Radially averaged cross-sections of the EDoF-Miniscope (solid red) and traditional miniscope (dashed blue) PSFs with a FWHM of 0.92 μm and 1.18 μm, respectively. (**H**) Raw measurements captured by EDoF-Miniscope of a neuronal population expressing GFP in a 100-μm thick fixed brain slice imaged through a cover glass (150 μm thick). (**I**) The same image after wavelet filtering. (**J**) The same region captured by a miniscope before (inset) and after wavelet filtering for comparison. (**K**) The maximum intensity projection (MIP) image of the same region acquired by a focal stack from a table-top widefield epi-fluorescence microscope with a 20x, 0.4 NA objective across a 100-μm axial range with a 10-μm step size.

We first evaluate the genetic algorithm designed DOE when integrating with a 1P miniscope. Our results show that the fabricated DOE closely matches the simulated profile and can be easily integrated into an EDoF-Miniscope with an alignment accuracy of 70 μm. We verify that the anticipated amount of lateral and axial misalignment of the DOE on the pupil plane negligibly affects the resulting EDoF. The resulting EDoF-Miniscope increases the DoF by ∼2.8x and achieves 0.9-μm lateral resolution when tested on 1-μm fluorescent beads. The DOE itself contributes 2 μg to the total weight and is directly etched on a 3.6 × 3.6 × 0.5 mm fused silica substrate, making it a compact solution for pupil engineering in miniscopes.

We experimentally demonstrate EDoF-Miniscope’s imaging capability across a variety of complex fluorescent samples. We show that EDoF-Miniscope successfully captures an extended fluorescent fibers volume in the presence of a strong out-of-focus background. We characterize EDoF-Miniscope on 5 μm and 10 μm fluorescent beads in scattering phantoms to demonstrate EDoF imaging in scattering media. We verify that the EDoF-Miniscope maintains a comparable SBR to a miniscope across scattering phantoms with different fluorescent bead densities. This proof of concept solidifies that the EDoF-Miniscope may successfully recover neuron-sized objects in scattering scenes. Lastly, we demonstrate the capability of EDoF-Miniscope in several brain samples. First, we image a fixed 100-μm thick mouse brain sample and compare the EDoF image to a widefield stack to show that we recover ∼82% of the total neurons within the volume. Next, we demonstrate the EDoF on imaging vasculatures in a fixed mouse brain.

Overall, our contribution is a novel EDoF-miniscope that achieves EDoF fluorescence imaging by integrating a lightweight DOE into a miniscope. We demonstrate the EDoF fluorescence imaging capability of EDoF-Miniscope across multiple fluorescent phantom and brain samples. Built from off-the-shelf components augmented by a customizable DOE, we expect that our low-cost EDoF-Miniscope may be adopted in many neuroscience labs and find utility in a wide range of neural recording applications.

## Results

### 2.1: Genetic Algorithm-based Design of DOEs on EDoF Miniscope

To design a DOE for deployment in 1P neural imaging, we develop a genetic algorithm. Our algorithm employs a linear shift invariant Fourier-optics model to optimize a binary DOE from a basis of three EDoF phase functions. In addition, we consider the native aberrations of the GRIN lens, as well as the exponential intensity decay due to tissue scattering. In this section, we highlight main insights into the algorithm design and refer to implementation details in **Methods 4.1, Supp. 1.1-1.3**.

A key feature of the EDoF-Miniscope image formation is the compression of axial information from an extended 3D volume to a 2D image. This requires that the optimized PSF must maintain a high contrast over an extended depth range to remain distinguishable from the out-of-focus background present in 1P neural imaging^1^. However, EDoF PSFs often achieve their “non-diffractive” properties at the expense of strong sidelobes that lowers signal contrast in dense scenes^29^. We address this issue by using a fitness function that judges the DOE based on how effectively it maintains peak intensity over the desired depth range, which reinforces the PSF to maintain a sharp contrast over the desired EDoF. In addition, we design the algorithm to optimize the EDoF PSF on neuron-sized objects (5-10 μm) by judging the resulting optical signal after convolving the 3D PSF with a simulated on-axis neuronal source at each depth. This allows the algorithm to further consider the geometric extension from finite-sized objects along with the diffractive effects from the binary DOE. We limit our analysis of the GRIN lens aberrations to third order on-axis seidel aberrations since higher order terms are dominated by high spatial frequencies on the pupil plane and the GRIN lens exhibits strong spherical aberration. At the end of each generation, our genetic algorithm refines the population by producing “children” masks that contain a mix of properties from the best masks in the current generation. By repeating this process over a few generations, the algorithm optimizes the desired EDoF pupil function. We analyze the convergence of the algorithm as a function of population size and number of generations in supp. We experimentally show that the resulting EDoF-Miniscope maintains a comparable SBR to a miniscope and that our designed PSF effectively elongates the PSF across the entire FoV for neural sized objects in **Supp. 5.1-5.3**

The input to the algorithm includes the scattering length (*l*_*s*_), refractive index (*n*), on-axis aberrations for the GRIN lens (*<u>W</u>*), number of pixels on our pupil plane (*N*) and properties describing our optical simulation (*<u>O</u>*) including the NA of the GRIN lens, system magnification, simulated FoV size, axial step size and number of depth planes for the simulated environment, and size of our proxy neuron on-axis source (see **Fig. 2A**). Within each “generation”, the algorithm optimizes over a set of basis phase terms, including axicon phase, defocus, and spherical aberration, to design a population of DOEs by solving the minimization problem, as illustrated in **Fig. 2B-C**

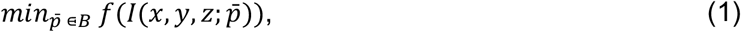

where 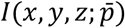 is the resulting 3D (*x, y, z*) intensity profile of the PSF parameterized by the pupil phase basis, 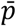, as determined by our forward model described in **Supp. 1.1**. *B* is the user-defined optimization bounds for 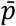. The fitness function *f* judges the quality of the EDoF PSF generated by each candidate within the population and takes the form:

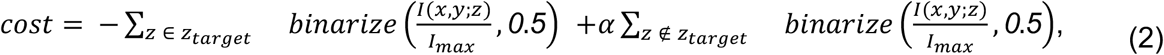

where z_target_ is the desired extension range, which is chosen based on the scattering length that sets the practical imaging depth limit for 1P fluorescence imaging. The operator *binarize*(*g, 0*.*5*) binarizes the 3D function *g* using 0.5 as the threshold, which effectively reinforces the desired EDoF PSF to retain at least 50% the maximum intensity I_max_ over the desired depth range across all depths. To prevent the algorithm from over extending the EDoF, we penalize any intensity profiles outside of the desired depth range by the *z* ∉ *z*_*target*_ term. *α*, which we heuristically set equal to 4, is a tunable parameter that determines the softness of the bound.

**Fig 2.**
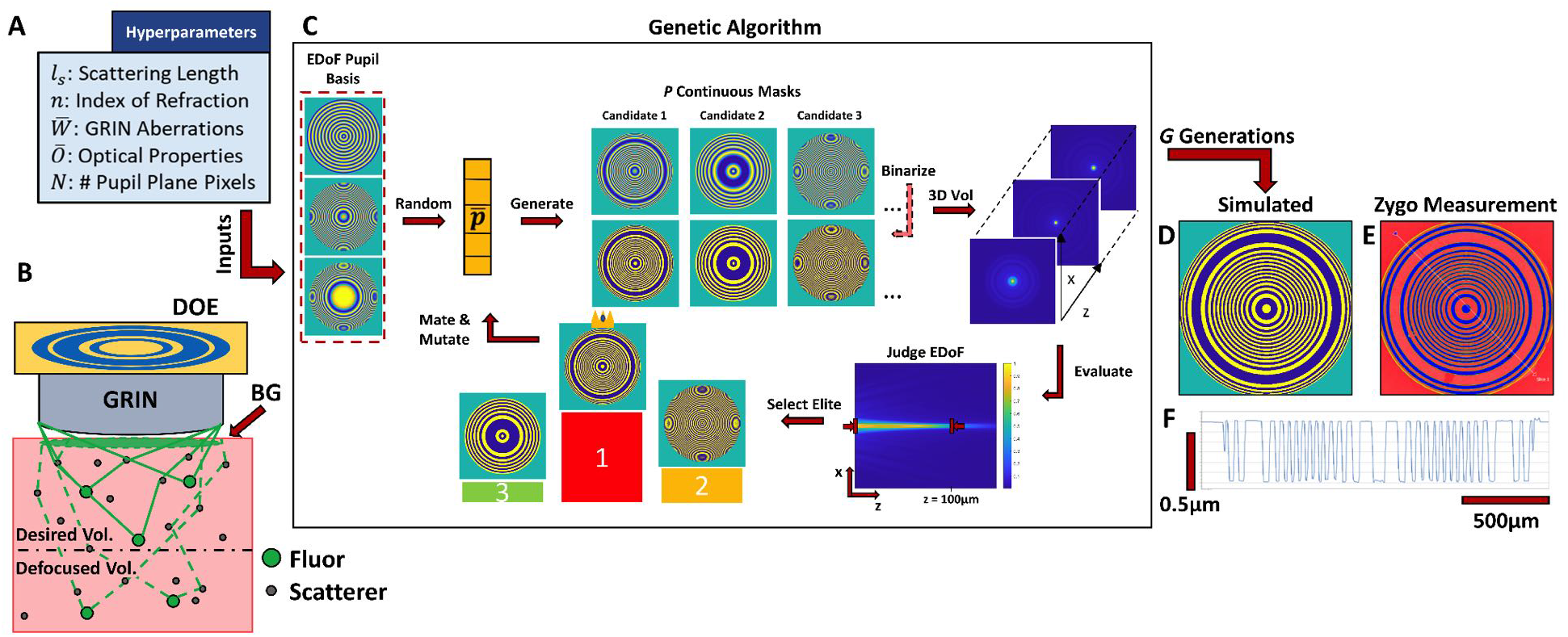
Genetic Algorithm for the Design of EDoF. (**A**) User-defined parameters used by the genetic algorithm to optimize the DOE. (**B**) Overview of the physical model used by the algorithm. We model the scattering effect by an intensity decay that scales with 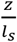. (**C**) Overview of the genetic algorithm process. First, the algorithm generates a population of candidate masks from a set of basis phase functions. These candidates are iteratively refined by judging their performance in extending the DoF within the scattering media. (**D**) Final simulated mask used for EDoF-Miniscope. (**E**) The final manufactured mask, and (**F**) its cross-section measured by a Zygo New View 9000 interferometer.

After optimization for a scattering length of 100 μm and background refractive index of 1.33, our binary mask is parameterized by a strong spherical aberration term, a strong defocus term and a negligible axicon contribution. By the binary nature of our mask, the +1st and -1st diffraction orders receive conjugate versions of the learned phase. It is important to note that each order will be subjected to the same aberrations, which may be modeled as a continuous pupil phase on top of our binary mask. In the +1st order, our mask adds an additional 26.65 waves of spherical aberration which exaggerates the native spherical aberration of the GRIN lens to greatly extend the focal range but behind the nominal focal plane. To compensate, the defocus displacement of 129 μm to bring the EDoF in front of the nominal focal plane. In the -1st order, the learned -26.65 waves of spherical aberration partially cancels the native aberration of the GRIN lens and displaces the focus by -129 μm to keep the order behind the nominal focal plane. As a result the +1st order is more extended than the -1st order. In addition the opposing defocus terms allow each order to produce non-overlapping axial profiles, so that our total DoF can be considered as the sum of each respective FWHM. Altogether our optimized binary mask is able to produce larger EDoF through our twin foci design, as shown in **Fig. 1D**. We further demonstrate our genetic algorithm’s flexibility to design DOEs to achieve an EDoF under different scattering conditions and background refractive indices in **Supp. 7**. Our genetic algorithm learns different weights of axicon, defocus and spherical aberration under each respective condition showing that the optimal binary mask varies as a function of the chosen physical parameters.

We select an optimized mask (**Fig. 2D-F**) and demonstrate its utility in imaging neural signals in brain tissues after being integrated into EDoF-Miniscope. We manufactured the mask using the single-step photolithography (see **Methods 4.3**) and aligned the DOE to the GRIN lens using the process described in **Supp. 3.1**. Practical considerations for integrating the mask, such as the effect of lateral and axial misalignment on the back pupil plane, are further studied in **Supp. 4.1**.

### 2.2: Experimental Demonstration on a Thick Fluorescent Fibers Sample

We experimentally verify the ability of EDoF-Miniscope on a *thick* complex fluorescence object by imaging fluorescent-stained fibers spread on a glass slide. The sample spans the full FoV (∼600 μm x 800 μm) and an extended depth (400 μm) to test the imaging performance of EDoF-Miniscope when subjected to a strong out-of-focus background. The EDoF-Miniscope raw measurement (see **Fig. 3A**, inlet) exhibits a lower contrast when compared to the miniscope raw measurement (see **Fig. 3B**, inlet), however this is due to the extended imaging range as well as an increased background. The SBR for the EDoF-Miniscope raw measurement is approximately 1.1 and 1.26 for the miniscope. As shown in **Fig. 3A**, the EDoF-Miniscope can clearly recover closely packed fiber structures across the full FoV after applying the wavelet filter to the raw data. The EDoF is highlighted by comparing the image to a miniscope (see **Fig. 3B**). Visually, the EDoF-Miniscope is able to recover fiber structures over sections of the FoV that are defocused in the miniscope image.

**Fig. 3.**
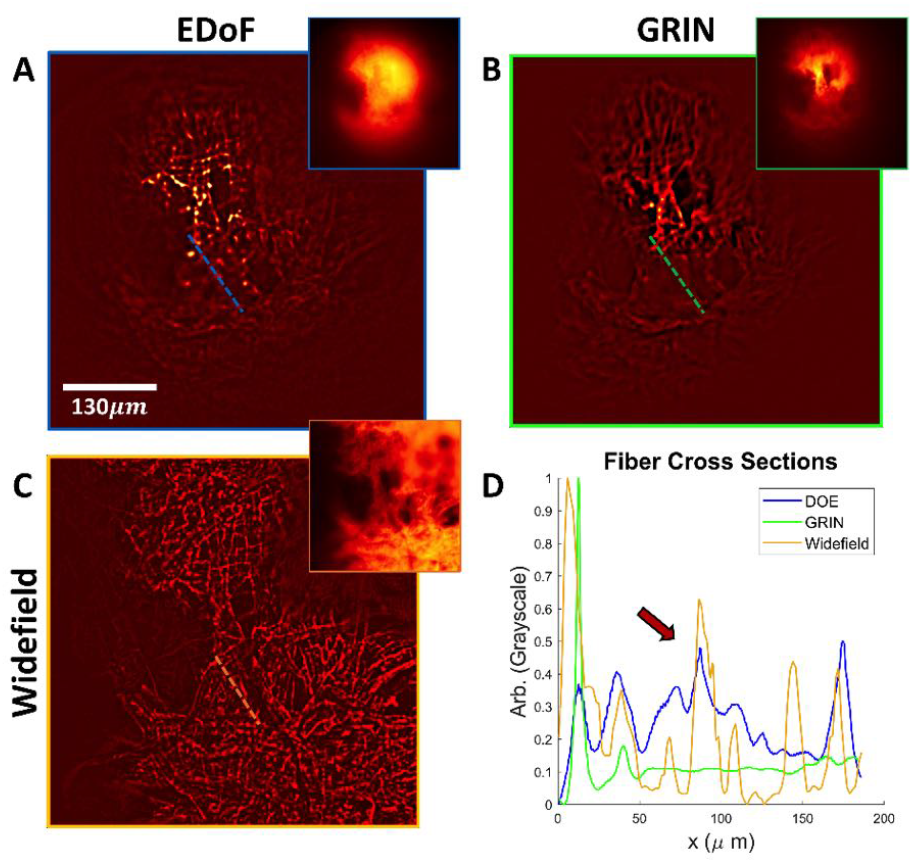
Extended Depth of Field in a Fluorescent Fiber Sample. (**A**) A fluorescent fiber sample imaged by the EDoF-Miniscope before (inlet) and after wavelet processing. (**B**) The same sample imaged by the miniscope and (**C**) MIP from widefield images collected over 125 μm with a 5 μm step size. (**D**) Cutlines across the same fiber bundle over the three previous images. The arrow indicates a region where the EDoF-Miniscope matches the synthesized widefield image but the miniscope is defocused.

To qualitatively show the EDoF-Miniscope’s EDoF capacity, we compare both images to an MIP obtained by projecting a z-stack of images collected by a tabletop widefield setup across 125 μm with a 5 μm step size (see **Fig. 3C**) using a 10x 0.25 objective lens (Nikon, CFI Plan Fluor, 10x, 0.25NA). We plot a cutline across a fiber bundle between each image, as shown in **Fig. 3D**. The EDoF-Miniscope retains more fibers than the miniscope when compared to the widefield MIP. The differences in the relative intensity scaling of the cutline between each image is due to the different illuminations used during the measurements (manually aligned epi-illumination for the miniscopes versus on-axis for the 10x objective lens) and aberrations. Visually, both miniscope images degrade towards the peripheries. Since the wavelet filter leverages subtle contrasts to extract features, its performance qualitatively degrades due to signal attenuation caused by severe off-axis aberrations as well. Notably the EDoF-Miniscope retains optical signals over a larger portion of the FoV than the miniscope, indicating that EDoF is partially resilient to off-axis aberrations. We further explore the off-axis properties of the EDoF-Miniscope in **Supp. 5.1-5.2**.

This experiment highlights the capability of the EDoF-Miniscope to extract fluorescent signals across the full FoV in the presence of an out-of-focus background. The complex and thick geometry of the fibers verifies that the EDoF-Miniscope may encode and extract arbitrarily shaped fluorescent sources through a simple wavelet filter to suppress any increased background. As a result, the EDoF-Miniscope can provide a high quality EDoF with good robustness to background signals.

### 2.3: EDoF in a Controlled Scattering Phantom

To further consider the EDoF-Miniscope towards neural imaging applications, we examine the performance of EDoF-Miniscope when encoding neuronal-sized (∼5-10 μm) sources under bulk scattering. To do so, we conduct experiments on a fluorescent bead phantom with scattering properties similar to that of neural tissue. Our phantom has a scattering length of ∼100 μm and an anisotropic factor of ∼0.97. We embed 5 μm fluorescent beads as proxy neurons in the scattering phantom to showcase the robustness of the EDoF-Miniscope to scattering media in this proof-of-concept experiment.

We image a spherical cap shaped phantom (diameter = 2.2 mm, depth = 0.65 mm) embedded with 5 μm fluorescent beads at a density of ∼2120 particles per mm^3^ (see **Methods 4.5**). Visually, the EDoF-Miniscope raw image (see. **Fig. 4A**, inlet) exhibits less contrast in the center of the FoV but retains more particles at the peripheries of the FoV when compared to the miniscope raw image (see. **Fig. 4B**, inlet). After performing a wavelet transform (see **Fig. 4A-B**) EDoF-Miniscope successfully extracts 111 particles versus 66 particles with the miniscope within the FoV. On average, the EDoF-Miniscope image exhibits an SBR of 1.06 while the miniscope exhibits an SBR of 1.10. We further explore the differences in SBR as a function of particle density in **Supp. 5.3**. and conclude that the EDoF-Miniscope trades on average a ∼4% decrease in SBR for its EDoF when imaging fluorescent bead samples (∼1000-10000 particles/mm^3^). We characterize the axial elongation of the 5 μm beads by both the EDoF-Miniscope and miniscope to judge the achievable imaging depth when interrogating non-diffraction limited, neuronal-sized sources. We accomplished this by axially sweeping a sparse scattering phantom with a custom automated sample stage (see **Supp. 3.2**). The EDoF-Miniscope achieves an imaging depth of 104 μm between both foci versus an imaging depth of 37 μm with the miniscope in **Supp. 5.1**. We designed the first order to have an elongation of 80 μm for 5 μm sized objects in scattering tissue; however here we observe an elongation of 67 μm. We predict that this discrepancy is due to the GRIN lens having a lower spherical aberration than what is predicted in our Zemax model. We repeat the procedure for 10 μm fluorescent beads and characterize the elongation on several locations across the FoV (see **Supp. 5.2**) to show that the EDoF-Miniscope also elongates off-axis sources, even if only on-axis aberrations are considered in our optimization.

**Fig. 4.**
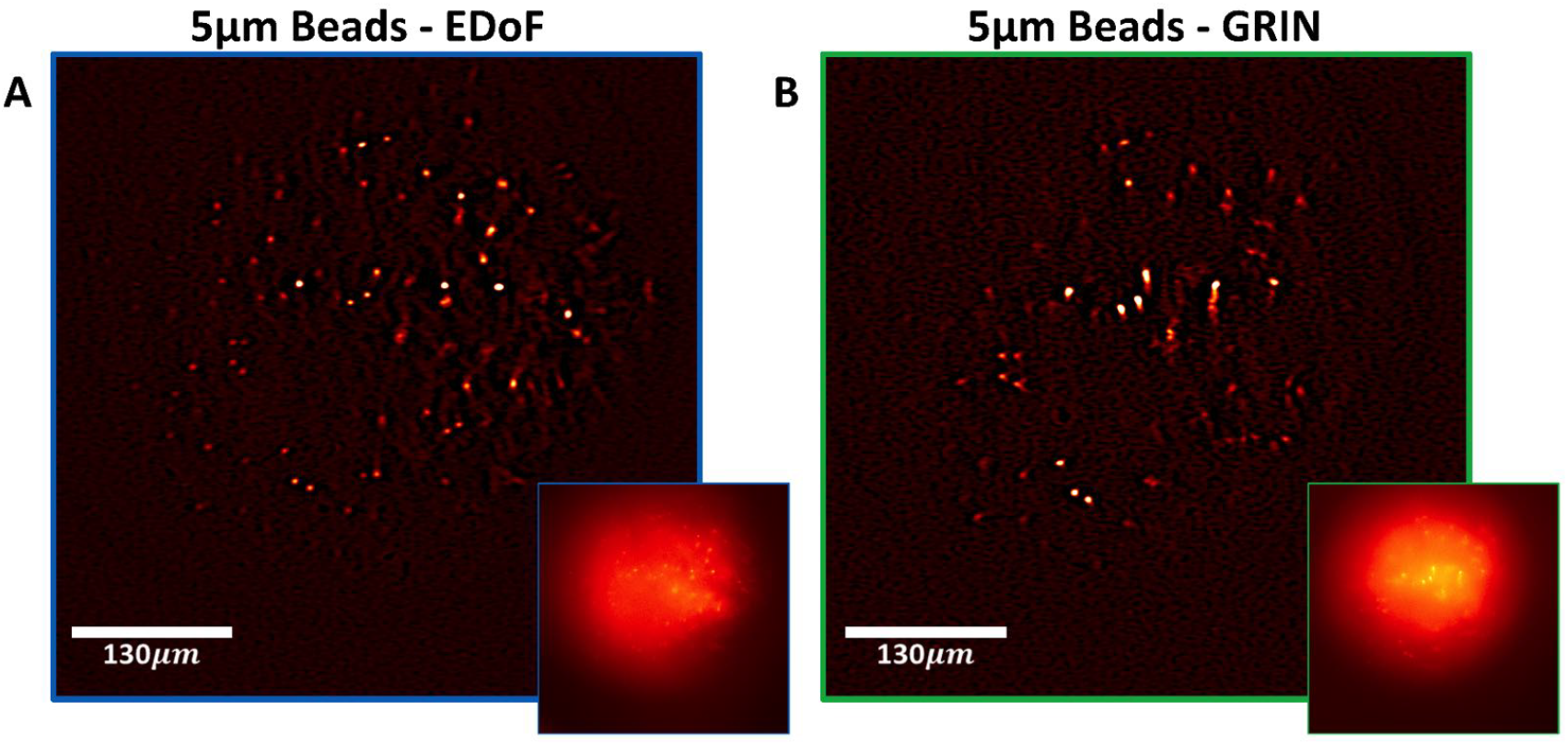
EDoF Characterization in Scattering Phantoms. (**A**) EDoF-Miniscope image of 5 μm beads embedded in clear resin with 1 μm polystyrene scatterers (ls = 100 μm) before (inlet) and after a wavelet filter. (**B**) The same region imaged by a miniscope, before (inlet) and after a wavelet filter.

### 2.4 Imaging Neuronal Structures in Brain Slices

To confirm the capacity of the EDoF-Miniscope to accurately extract neuronal structures, we image the same neuronal population of a 100 μm thick fixed brain slice stained with GFP in an axial sweep with a widefield tabletop imaging system (see MIP in **Fig. 5A**) and in a single-shot with an EDoF-Miniscope with (see **Fig. 5B**) and without wavelet filtering (see **Fig. 5B**, inlet). The widefield stack was acquired on a sCMOS camera (PCO.Edge 5.5, pixel size = 6.5 μm) with a Nikon 20x 0.4NA objective by performing a z-scan 100 μm depth range with a 10 μm step size. Visually, both the EDoF-Miniscope image and widefield MIP capture neuronal structures in the center of the FoV, however, the MIP has better performance near the peripherals due to the less extreme off-axis aberrations in the objective lens. Despite this disparity, the EDoF-Miniscope recovers 174 neurons in a single shot and the widefield MIP recovers 213 neurons over its z-scan range.

**Fig 5.**
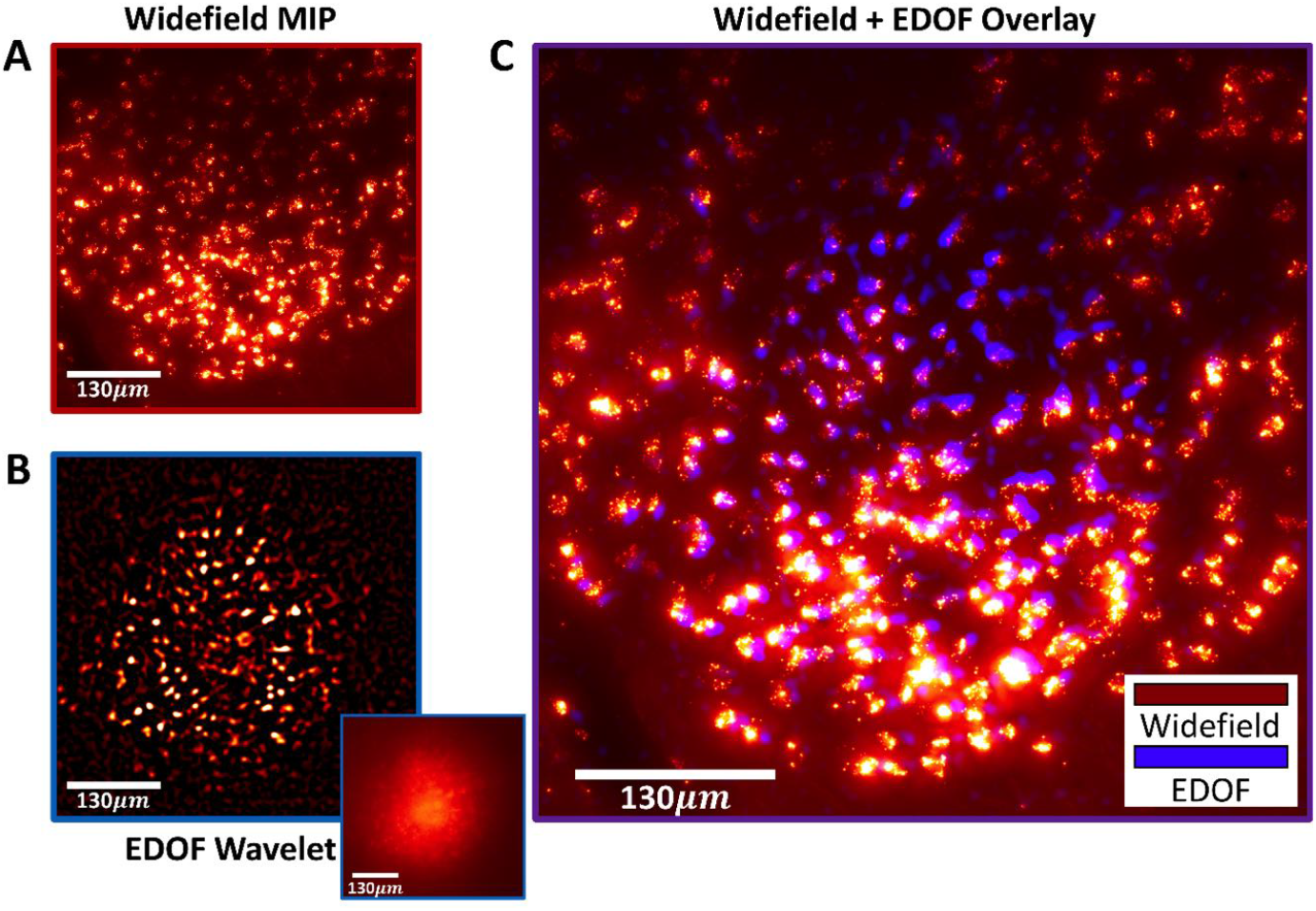
Confirming Wavelet Extraction of Neuronal Structures. (**A**) Widefield MIP of the thin brain slice captured over a 100 μm depth range with a 10 μm step size. (**B**) singleshot EDoF-Miniscope image of the same neuronal population. (**C**) Overlaid EDoF-Miniscope and Widefield frames after performing resampling on the widefield frame to place them on the same spatial grid.

Next, we overlay the images to perform additional visual inspections. We resample the widefield MIP to the same discretization as the EDoF-Minsicope image and crop the images to the same FoV. We confirm that the structures extracted in the wavelet processed EDoF-Miniscope frame match well with the neuronal structures recorded in the widefield MIP. This result confirms that our framework can properly extract neurons from the background without artifacts. It is important to note that the widefield MIP and EDoF-Miniscope images are not in perfect agreement across the whole FoV. This variation is due to misalignment in the miniaturized imaging system combined with distortions in the miniaturized optics that are not accounted for in the overlaying process. However, these factors only affect the neuronal positions mildly and visual verification is still possible across the FoV. We additionally visually verify that the wavelet filter performs superior neuronal extraction than direct deconvolution with morphological background removal for the EDoF-Miniscope image in **Supp. 6.2**

### 2.5: EDoF Imaging of Vasculatures in a Fixed Whole Mouse Brain

We imaged fluorescently stained vessels in a fixed whole mouse brain sample. As shown in **Fig. 6A-B**, we imaged the same vasculatures under both the EDoF-Miniscope and a miniscope such that the pronged vessel in the center of the FoV was in focus. This central vessel exhibits an SBR 1.09 of for the EDoF-Miniscope and 1.11 for the miniscope. Visually, both raw measurements are sparse allowing us to directly inspect the structures. Applying the wavelet filter allows us to interrogate the fine features and inspect the relative sizes of the vessels. A further comparison of wavelet processing versus deconvolution and morphological background removal can be found in **Supp. 6.3**. The miniscope image retains fewer capillaries and exhibits a broadening of the large vessel when compared to the EDoF-Miniscope image. This result highlights that the EDoF-Miniscope is able to capture diverse types of fluorescent objects.

**Fig. 6.**
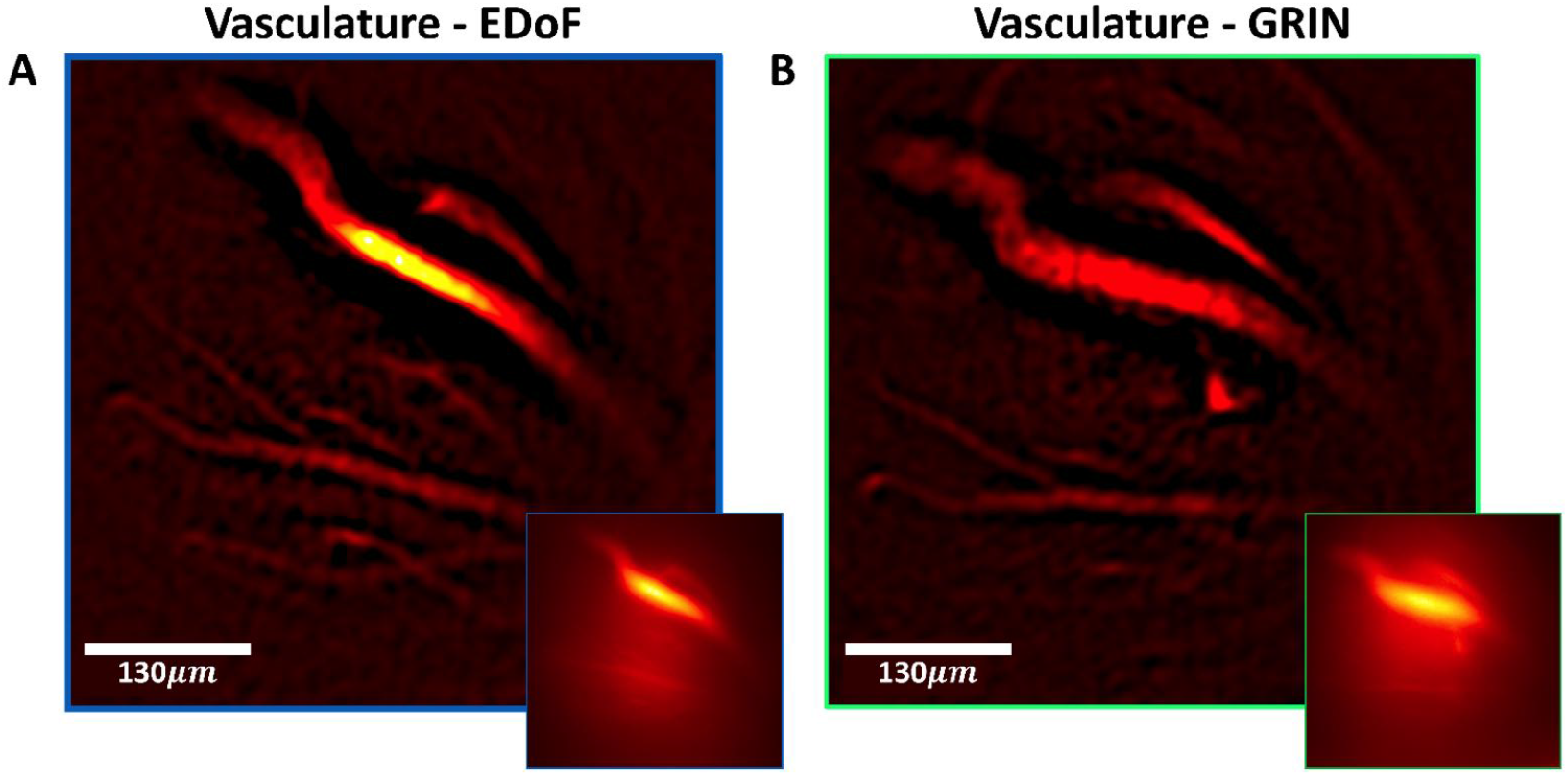
EDoF Imaging of vessels in a Fixed Mouse Brain. (**A**) Fluorescently stained vessels in a fixed whole mouse brain captured by a EDoF-Miniscope. The EDoF-Miniscope was placed such that the pronged vessel was centered in the DoF. (**B**) The same region captured such that the pronged vessel was in-focus on a miniscope.

## Discussion and Conclusions

In summary, we have presented a novel EDoF-Miniscope, which applies a binary phase DOE to augment the miniscope for EDoF imaging. The system uses a genetic algorithm to design an EDoF for deployment in 1P fluorescent imaging and achieves a ∼2.8x improvement in DoF when compared to a miniscope. We have experimentally verified the EDoF-Miniscope across a variety of complex fluorescent and neuronal structures.

The main contributions of the EDoF-Miniscope is its novel design and integration of binary DOEs into miniscopes to achieve single-shot recording of neural signals across an EDoF without sacrificing spatial resolution nor optical multiplexing. The genetic algorithm synthesizes a DOE by refining a pool of candidates from a basis of three EDoF phase functions to deploy a scattering-robust EDoF for neural imaging. After manufacturing, the DOE itself weighs only ∼2 μg and may be integrated into an EDoF-Miniscope with an accuracy of 70 μm without degraded performance. The EDoF-Miniscope exhibits a slightly decreased SBR to a miniscope. We use a simple wavelet filter to extract extended fluorescent signals across the full FoV.

Our pilot demonstration on the utility of DOEs in miniscopes may be a particularly attractive area for future research as it exploits the open source miniscope project that encourages customizability and rapid development for usage in a diversity of experiments. By promoting an easily manufacturable and customizable DOE and requiring minimal modification to the miniscope architecture, the EDoF-Miniscope may offer a customizable framework that allows researchers to better tailor miniscopes to match their experimental requirements. Broadly, we expect that the synergy between physics-based computational design strategies and customized miniature phase masks will continue to improve interrogation of neural signals in miniaturized microscopes and endoscopes with additional novel capabilities, such as extended FoV^30^.

In its current prototype, the EDoF-Miniscope is designed for performing proof of concept experiments in fixed samples in a tabletop setup. In future applications, we intend to replace our 230 μm working distance GRIN lens with a 0 working distance GRIN lens, which is more suitable for in-vivo studies^3^. We will also utilize a backside illuminated (BSI) CMOS sensor to significantly improve the SNR and dynamic range for in-vivo studies. While cutting edge sensors may increase the size and weight of the miniscope^11^, we are encouraged by the recent development of the MiniFAST BSI CMOS-based miniscope^31^ in successfully applying high data rate and high pixel count BSI sensors to future generations of the EDoF-Miniscope. Our studies indicate that we may further decrease the size and weight of the platform while nominally affecting its imaging properties to further reduce the formfactor (see **Supp.1.2**) .

An outstanding challenge in expanding the EDoF-Miniscope towards in-vivo studies is overcoming the high background present in in-vivo neural imaging. There are several promising solutions we envision for future generations of the EDoF-Miniscope, such as replacing the binary DOE with a miniature refractive element^8,30^, incorporating structured illumination techniques^5,32^, and advanced computational techniques to facilitate neuronal signal extraction^33–35^.

## Methods

### 4.1: Genetic Algorithm Design and Implementation

We developed the genetic algorithm using the Genetic Algorithm Toolkit in Matlab 2019b to optimize the DOE over a forward model to minimize a fitness function that encourages EDoF behavior. First, we define a target number of generations, *G*, for the algorithm to optimize over and select the population size, *P*, per generation. Next, we define several optical and physical parameters. For the DOE demonstrated in the EDoF-Miniscope, we used a scattering length (*l*_*s*_ = 100 μm), refractive index (*n* = 1.33), on-axis (spherical) aberrations for the GRIN lens (*W* = *W*_*040*_ = 29.4), number of pixels on our pupil plane (*N* = 1000 × 1000) and properties describing our optical simulation (*O*) including the NA of the GRIN lens (0.55), system magnification (9.2x), real space pixel size (3.45 μm) axial step size (1 μm) and number of depth planes for the simulated environment (100), and size of our proxy neuron on-axis source (5 μm) to reach a target depth of 80 μm in the first diffraction order. During a single iteration, the genetic algorithm selects a candidate basis, *x*, from the current population of *N* bases and generates the binary phase for our DOE using:

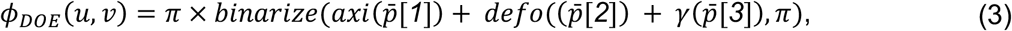

where *axi* is an axicon phase kernel described by 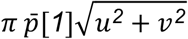, *defo* is an angular spectrum defocus phase kernel described by 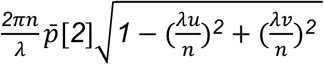, *γ* is a spherical aberration phase kernel described by 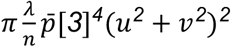, and *(u,v)* are spatial frequency coordinates on the pupil plane. The resulting phase is wrapped and binarized to 0 and *π* to achieve our binary phase. Next, we generate our resulting pupil plane profile by:

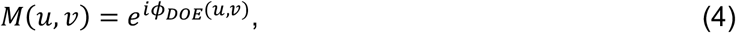

where *M(u,v)* is a candidate binary filter for use on the pupil plane of the miniscope. Since our binary phase assumes the values 0 or *π*, our resulting DOE phase factor will correspondingly assume the values 1 and -1 respectively. Next, the algorithm simulates the slice-wise 3D intensity distribution over the desired depth range by:

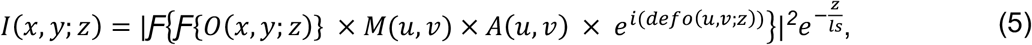

where *I(x,y;z)* is a 3D intensity distribution, *O(x,y;z)* is an object function representing a target-sized neuron placed on the optical axis, *A(u,v)* is the on-axis aberrations from the GRIN lens modeled on the pupil plane, and the last term accounts for the expected intensity loss due to scattering while neglecting the negligible amount of PSF width broadening^36^ within a single scattering mean free path. As a result, each slice in *I(x,y;z)* represents the optical signal generated by a neuron placed at a distance *z* in the expected environment. By limiting our simulation to on-axis aberrations, we may utilize a fast and efficient linear shift invariant model, which improves the convergence speed of the algorithm.

After the algorithm determines the cost for each candidate in the current generation, it develops three subpopulations to transfer to the next generation, including the elite children, crossover children, and mutation children. Elite children are replicas of the best performing candidates within a top percentage (in our case 20%) of all fitness values within the current generation, allowing these well-performing traits to transfer unperturbed to the next generation. Crossover children are non-elite children generated by randomly selecting a subset of the population (in our case 4 candidates) and combining the traits of the two best masks within that population. Mutation children are non-elite children generated by the same process as crossover children, however the traits of each child undergoes a random chance to mutate after creation. In our algorithm, each trait undergoes a 10% uniform chance of mutation, where mutation replaces the current value with an argument selected from the allotted bounds. The role of elite children is to keep the best traits unperturbed between generations to enforce that the next generation must contain a fitness value as good as the previous generation. This strategy is known as a “queen bee” genetic algorithm and guarantees global convergence after enough time^37^. Conversely, crossover children and mutation children are designed to reshuffle the combination of traits in circulation in an effort to discover new optimums *without* the induced bias of the elite candidates. The crossover fraction (in our case 0.4) determines the fraction of the non-elite children that will be generated through crossover instead of mutation. Once the genetic algorithm selects an optimal mask, we determine the sensitivity of the mask due to lateral displacement and axial displacement on the pupil plane as shown in **Supp. 4.1**. For 10 generations with 60 masks per generation, the algorithm typically converges in 80 minutes. Additional details about the genetic algorithm convergence and completion time are in **Supp. 2**.

### 4.2: Design and Characterization of EDoF-Miniscope

The EDoF-Miniscope is a standalone miniature fluorescence microscope that is built with off-the-shelf optical components, a custom DOE and a 3D-printed housing, as illustrated in **Fig. 1A-B**. The EDoF-Miniscope consists of an epi-fluorescent architecture consisting of an illumination and imaging path to excite and collect fluorescent signals within a circular field of view of ∼800 μm in diameter. The full breakdown for the EDoF-Miniscope is in **Fig. 1A**. For the illumination path, a surface-mounted LEDs (Luxeon, Rebel Blue) is aligned to a 3D printed epi-fluorescent channel in the miniscope. The LED is connected to a driver (LED dynamics Inc., 3021-D-E-350, 350A). The LED illumination first passes through a ball lens (Edmund Optics, 45-549) and is spectrally filtered by the excitation filter (Chroma, bandpass filter, 480/40 nm, 4mm × 4mm × 1.05mm). The filtered illumination reflects off a dichroic mirror (Chroma, 500 BS, 4mm × 4.8mm x 1mm) and is focused into the sample through the modified GRIN lens. Since the LED can be considered spatially incoherent, we can ignore the effects of the DOE on the illumination beam. For the imaging path, we relay the generated fluorescent signal through the modified GRIN lens and spectrally filter the fluorescent signal with an emission filter (Chroma, bandpass filter, 535/50 nm, 4mm × 4mm × 1.05mm). The filtered signal is magnified 9.2x and focused by an achromatic lens (Edmund Optics, NT45-207, f = 15 mm) onto an external monochrome CMOS camera (FLIR, BFLY-PGE-50A2M-CS). We modeled the imaging path in Zemax to determine the optimal placement of the optics (see **Supp. 1.2**).

To integrate the DOE into the EDoF-Miniscope, we glue (Noland Products Inc., NOA63) the DOE to the back surface of the GRIN objective lens (Edmund Optics, 64-520) with a 70 μm precision to produce the modified GRIN lens. Next, we place the modified GRIN lens into a 3D printed objective lens holder and adhere it onto our miniscope body by curing dental paste (Pentron, Flow-It ALC, Opaque A1) with a UV source (Alonefire, SV003, 10 W 365 nm UV Flashlight). The thickness of the DOE substrate (500 μm) is close to the 230 μm working distance of the GRIN lens allowing us to approximate that the glued DOE is on the back focal plane of the GRIN lens. We confirmed that the lateral and axial placement of our DOE is within our tolerances in **Supp** .**4.1**. As compared with a miniscope, the assembled EDoF-Miniscope achieves a ∼2.8x improvement in DoF without sacrificing optical resolution.

The EDoF-Miniscope is comparable in size and weight to a standard GRIN-based miniscope, and incorporates additional features, and is demonstrated on a tabletop setup for proof-of-concept (see **Supp 3.2**) with an automated z-sample stage. The top plate is designed to block light leakage from the on-board LED into the external camera and side fin allows the EDoF-Miniscope to attach to an O1/2” Thorlabs (Thorlabs, TR2) post to attach to a linear 1” XYZ stage (Thorlabs, PT3A). We integrate the modified GRIN lens into an EDoF-Miniscope using a 3D printed objective mount, as shown in **Fig. 1A-B**. With modifications for tabletop use, the resulting EDoF-Miniscope weighs 2.3g and has a size of 22.39mm x 16.5mm x 15.6mm. Without these modifications, the platform weighs 1.85 g and has a size of 13.17mm x 8mm x 15.6mm.

We compare the PSF of EDoF-Miniscope to a miniscope by scanning a 1 μm fluorescent bead (Thermo Fisher Scientific, Fluoro-Max Dyed Green Dry Fluorescent Particles) on a glass slide along *z* to characterize the DoF of each platform respectively using the automated test setup. As seen in **Fig. 1C**, the EDoF-Miniscope produces asymmetric imaging foci around the nominal focal plane, as predicted by our Fourier optics-based model. By radially averaging the maximum intensity projection (MIP) of the EDoF-Miniscope and miniscope 3D PSFs over their focal range, we may compare their average optical resolution over their imaging range. The EDoF-Miniscope produces a comparable resolution to the miniscope, indicating that our optimized PSF does not trade its lateral resolution for the EDoF. The slight differences in the measured resolution is likely due to the manual alignment of the optical components within the EDoF-Miniscope and miniscope, respectively. Noticeably, the EDoF-Miniscope PSF is characterized by a strong defocus lateral ring, which enables the EDoF behavior and contributes to an increase in background. We can see this increased background by comparing the raw measurement of the EDoF-Miniscope on a 100 μm mouse brain slice (see **Fig. 1H**) to the raw measurement from a Miniscope (see **Fig. 1J** inlet). However, **Fig. 1I-J** shows that we recover more signals with the EDoF-Miniscope than the miniscope after applying a simple wavelet filter. This experiment indicates that the DOE is effective at encoding neural signals across an extended depth.

### 4.3: Fabrication of the binary Diffractive Optical Element

We manufactured the binary diffractive optical element using single step photolithography in the Boston University Class 100 Optoelectronics Processing Facility cleanroom. We first develop a .dxf file containing a 3×5 array of optimized DOEs within a 11.4mm x 19 mm area using AutoCAD and convert the file into a .gdsii using LinkCAD. We upload the .gdsii file onto an optical mask writer (Heidelberg, DWL66) to write the DOE pattern into the photoresist layer of an optical mask blank (Nanofilm, 5×5X.090-SL-LRC-10M-1518-5K). Next, the exposed optical mask is placed in photoresist developer (Microposit, Photoresist Developer MF-319) followed by chromium etchant (Transene, Chromium Etchant Type 1020) for 1 min, which exposes a transparent soda lime glass corresponding to the phase shifted rings in each DOE. The remaining photoresist is removed by submerging the optical mask in a bath of 80°C photoresist remover (Microposit, Photoresist Remover 1165) for 20 min. We next prepare to develop the masks onto a 50.8 mm diameter by 0.500 mm thick fused silica wafer (University Wafer #971) by first singling the substrate in a convection oven at 120 °C for 15 min. We spin photoresist (Microposit, S1813) onto the wafer at 4000 rpm using a photoresist spinner (Headway Research PWM32-PS-CB15) for 45 sec and soft bake the resist at 90°C for 15 min. Next the wafer and optical mask are aligned using an optical mask aligner (Karl Suss, MA6) and the wafer is exposed the 10 mW/cm^2^ *λ* = 365 nm UV source onboard for 16.4 sec, or until a dose of approximately 150 mJ/cm^2^ was achieved. We added 50 μm of separation between the chrome mask and our wafer to match our desired resolution of 7 μm in accordance with the contact printing equation^38^:

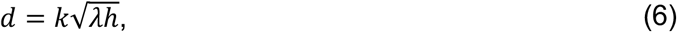

where d is the resolution of the photolithography procedure, k is a constant which is 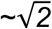 for photolithography, *λ* is the wavelength of light and h is the thickness of the photoresist. After exposure, we hardbake the wafer at 120 °C for 10 min. Next, we develop the mask in a diluted solution of MF-319 and DI water [2:1] for 75 sec to reduce dark erosion and improve the mask quality. Once the photoresist is developed, we etch the final pattern in a reactive ion etcher (Plasma-Therm, Reactive Ion Etcher) using a mixture of CHF_3_ and O_2_ (45-5 sccm) at 40 mTorr pressure for 16.8 min or until the phase shifted rings reach their target depth in accordance with the target phase shift. The array of DOEs are then aligned to a cutting blade (Disco, DPLU0921) and diced using an automated dicing saw (Disco, DAD 3220). Any remaining photoresist is stripped away in an 80 °C bath of 1165 remover. We measure the etched depth in an optical interferometer (Zygo, New View 9000) to justify the etch quality, as shown in **Fig. 2D-E**.

### 4.4: Quantification of Extended Depth Range

To quantify the DoF improvement in experimental results, we use an automated stage to scan a target object across z while being imaged by the EDoF-Miniscope or a miniscope. Next, we open the resulting xy image stack in Fiji ImageJ and reslice the array into a stack of xz perspectives. We perform maximum intensity projection on the xz stack to visualize the 3D optical signal. We extract a line profile detailing the optical signal along the optical axis. We define the recoverable depth range as the Full-Width-Half-Maximum of the optical signal. In practice, this quantity represents the maximum depth range we may observe before the optical signal is reasonably obscured by the high background commonly observed in 1-photon neural imaging.

### 4.5: Scattering Phantom Preparation

We prepared the scattering phantom samples by following the procedure presented in our prior work^11^. In brief, we embedded 5 μm fluorescent microspheres and a separate sample of 10 μm microspheres (Thermo Fisher Scientific, Fluoro-Max Dyed Green Dry Fluorescent Particles) in a background medium of clear resin (Formlabs, no. RS-F2-GPCL-04; refractive index is approximately 1.5403) to act as idealized neurons. We controlled the bulk scattering of the phantom by embedding 1.1 μm nonfluorescent polystyrene microspheres (scatterers) (Thermo Fisher Scientific, 5000 Series Polymer Particle Suspension; refractive index, 1.5979) at a controlled amount. We estimated the effective ls of the phantom through the use of Mie scattering theory:

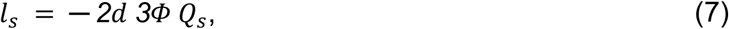

where *d* is the mean diameter of the scatterers and *Φ* is the volume fraction of the scatterers. The scattering efficiency, *Q*_*s*_, is derived with a Mie scattering calculator. To match a scattering length of neural tissue (100 μm) we add 0.27 mLof a scattering suspension (10% by volume) per mL of clear resin into the phantom. After mixing the suspension with the resin, we leave the resin in a dark location overnight. This period allows the water from the suspension to evaporate away without curing the resin and ensure the background medium remains homogeneous, except for the contribution of the scatterers. The resin is then moved to a glass slide and cured under a high power UV torch for 30 seconds.

### 4.6: Whole Fixed Mouse Brain Sample

We prepared the *ex-vivo* whole brain sample by preparing a mixture of gelatin (Sigma Aldritch, G2500) in PBS (ThermoFisher Scientific, 10x pH 7.4, 70011069) diluted 1:10 at 10% weight-per-volume for a maximum amount of 10ml for one animal. The mixture was placed on a hotplate and heated to 40-45°C to keep the gelatin dissolved. Next a heparinized PBS mixture is prepared by adding 600 iu of heparin (ThermoFisher Scientific, H7482) to 30 ml of PBS and is kept at 40-45 °C. Next, 30 mg of FITC-albumin (ThermoFisher Scientific, A23015) is added to 1 ml of PBS. Just before perfusion, we filled a beaker with crushed ice and added the FITC-albumin solution to the gelatin solution, shaking gently. Next we performed a cardiac perfusion with the heparinized saline followed by the FITC-albumin-gelatin. The mouse was put head down into the crushed ice for 15 minutes. Afterwards the brain was extracted, placed in 4% PFA (Electron Microscope Sciences, Diluted 1:8, 15714S) for 6 hours then placed in PBS. To finish the fixation process, the brain in PBS was placed on a horizontal shaker for 3 days.

### 4.7: Zemax Simulation

We conducted a series of Zemax simulations to study the imaging path of the EDoF-Miniscope. We used a sequential mode simulation to analyze how the PSF changes as we adjust the placement of the filters and lenses in the EDoF-Miniscope body. We judged the performance of a certain configuration by setting an RMS spot size merit function at the camera plane when imaging an on-axis and in-focus diffraction limited spot at our target wavelength. To optimize our design, we performed hammer optimization while constraining the design space so that the total lens distances did not exceed the maximum allotted size for the EDoF-Miniscope. We analyze how the location of the optics affects the PSF size. We also use our simulation to determine the expected on-axis aberrations generated by the EDoF-Miniscope. This process allowed us to optimize the performance of the EDoF-Miniscope under our experimental condition and inform our genetic algorithm of the aberrations we need to optimize over. We modeled the geometric effects of the phase mask substrate by adding a 500 μm thick layer of fused silica at the back surface of the GRIN lens.

### 4.8: Automated Data Collection through Pycro-Manager

We created an automatic data collection pipeline by automating the motion of a Thorlabs single-axis stage (Thorlabs, PT3A) and our sCMOS acquisition using Pycro-Manager^39^. In brief, Pycro-Manager allows for the seamless control of popular microscopy components, cameras using python routines in an easy to use framework that builds on Micromanager’s core functionality. We captured our diffraction limited PSF (see **Fig. 1D**) by bringing a dried 1 μm fluorescent bead (Thermo Fisher Scientific, Fluoro-Max Aqueous Green Fluorescent Particles) on a glass slide into focus using our experimental setup. Next we used our Pycro-Manager stage to scan the sample in z with a 1 μm step size from -150 μm to 150 μm. We set the exposure time to maximize the in-focus signal without saturation using the camera with 16-bit discretization, however the framerate never fell below 30Hz. We repeated the procedure for our 5 μm and 10 μm bead scattering sample with a 1 μm and 5 μm axial step size, respectively. We used Pycro-Manager to automatically apply an optimized wavelet filter to each captured frame to save both a raw and wavelet processed stack during an acquisition.

### 4.9: Wavelet Filter Design and Implementation

Our wavelet closely follows a wavelet filter proposed for extracting fluorescent signals from a low contrast measurement^25^, which takes the form:

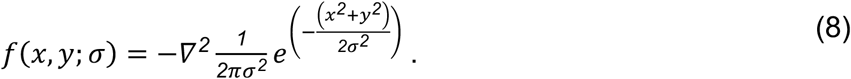

This kernel, known as the Laplacian of Gaussian (LoG), reduces noise sensitivity in a measurement while highlighting rapid changes in intensity in a single, easily precalculated kernel. While commonly used in edge detection^40^, prior work^25^ showed that adapting the standard deviation of the kernel to the approximate size of the fluorescent object provides targeted background suppression while reducing noise. We confirm the properties of the LoG kernel in **Supp. 6.1**.. We apply the wavelet to our simulated data using Matlab and to our experimentally collected data in Python. In both instances, we determine the optimal *σ*, which is commonly 8 pixels, and preallocate the fourier transform of the target LoG kernel. Afterwards, we may rapidly apply the kernel in Fourier space to clean the target measurement using minimal computation resources.

## Code availability

The genetic algorithm implementation and post-processing algorithms are available at: https://github.com/bu-cisl/EDoF-Minsicope

## Acknowledgements

The authors acknowledge funding from the National Institutes of Health (R01NS126596). Joseph Greene acknowledges the National Science Foundation Neurophotonics Research Trainee: Understanding the Brain: Neurophotonics Fellowship (DGE-1633516). The authors thank Boston University Optoelectronics Processing Facility for providing the resources and training to perform photolithography, the Boston University Precision Measurement Lab for access to the Zygo NewView 6000 optical profilometer and Boston University Photonics Center for providing access to Zemax.

## Author contributions

J.G. and L.T. conceived the idea. J.G. manufactured the diffractive optics, prototyped the hardware platform and algorithm, and conducted the experiments. Y.X. assisted with designing the platform, and conducting and analyzing experiments. J.A. helped with assembling the system, preparing experiments and analyzing data. A.M. helped design the alignment setup and develop the forward model. G.H. helped with the experiments. K.K. and I.D. prepared the biological samples. All authors participated in writing the paper.

