## supplemental material for "EDoF-Miniscope: pupil engineering for extended depth-of-field imaging in a fluorescence miniscope"

#### Contents


1 Optical Modeling & Characterization 2
1.1 Forward Model Implementation 2
1.2 System Characterization in Zemax 3

2 Genetic Algorithm Convergence 5

3 Optical Setups 6
3.1 High Precision DOE Alignment Setup 6
3.2 Automated Test Platform for Fixed Samples 7

4 DOE Perturbation Analysis 8
4.1 **Effect of Lateral and Axial DOE Displacement on the DoF** 8
4.2 Alignment Precision under Visual Guidance 8

5 EDoF Characterization on Fluorescent Samples 10
5.1 5-μm Fluorescent Beads 10
5.2 10-μm Fluorescent Beads 11
5.3 SBR Analysis Over 5-μm Bead Phantoms of Varying Density 12

6 Wavelet Transform Analysis 13
6.1 Simulated Wavelet Performance 13
6.2 Comparison to Deconvolution - Brain Slice Sample 14
6.3 Comparison to Deconvolution - Vasculature 15

7 Exploration of DOE Optimization in Different Scattering Media 16

8 Citations 17

### 1 Optical Modeling and Characterization

#### 1.1 Forward Model Implementation

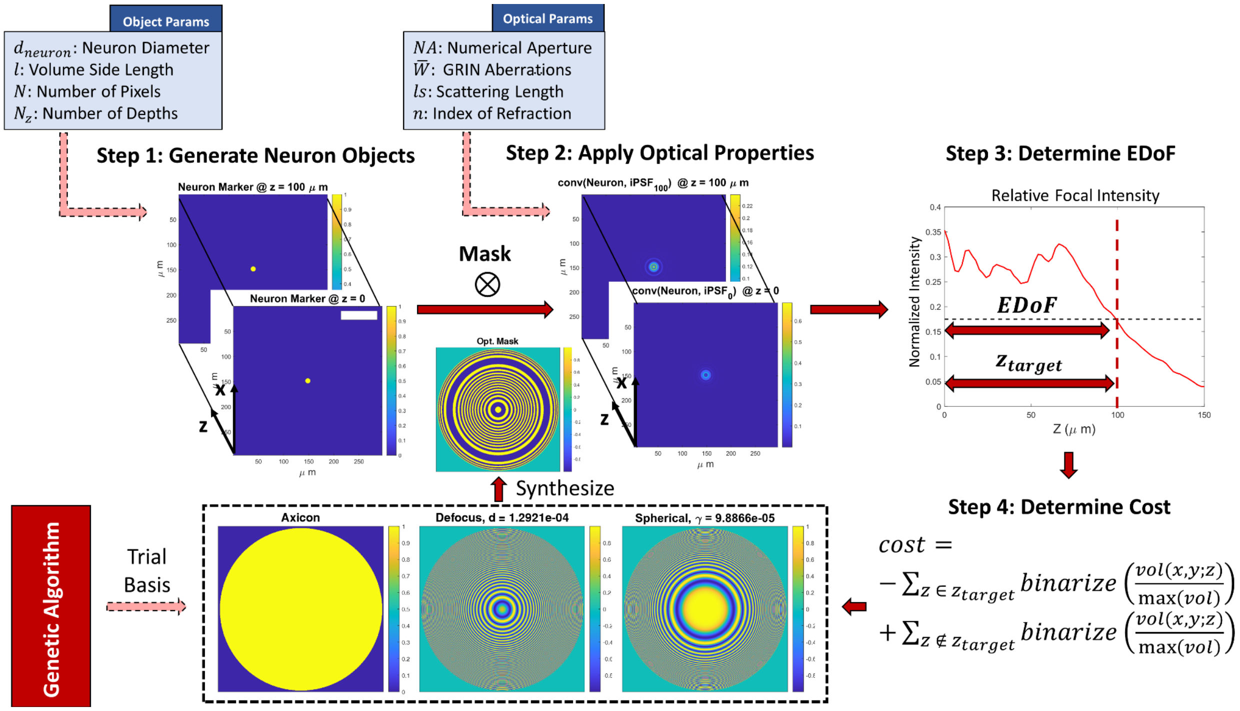


**Fig S1.1. Genetic algorithm forward model.** The genetic algorithm selects a candidate and generates the 3D optical signal for a neuronal sized object. The forward model leverages user-defined information to generate the appropriate scene. After generating the 3D signal, our algorithm judges the EDoF on the full-width-half-maximum of the on-axis XZ cross-section. Applying the fitness function to each candidate allows the algorithm to refine the current population and produce children for the next generation.

The imaging model used in this work follows a scalar diffraction approximation to describe the light emitted from an isotropic fluorescent emitter [Goodman]. Our optical platform is a 4f system with a diffractive optical element placed on the pupil plane. We assume that the emitters are suspended in a user-defined media described by an expected refractive index, $n_m$, and a scattering length, $l_s$. We assume that the media is strongly forward scattering, as is common with biological tissue[1]. Under these assumptions, we may characterize the optical signal resulting from an on-axis neuronal source on our image plane by:

$$I(x, y; z) \;=\; |F\{F\{o(x, y; z)\} \times M(u, v) \times D(u, v; z) \times A(u, v)\}|^2 \times e^{-z/l_s}, \tag{S1}$$

where *(x,y)* is our magnified spatial coordinates, *z* is the current depth relative to the focal plane, and *(u,v)* are the corresponding spatial frequencies. $o(x, y; z)$ is our on-axis object function, which we typically assume is a 5 µm circle as proxy neurons. To characterize the EDoF, we simulate on-axis emitters where the thickness of our fluorescent sources matches our axial resolution. The volume is described by hyperparameters set by the user before running the simulation (see **Methods 4.1**). $M(u, v)$ is the field imparted by our simulated binary mask placed on the pupil plane and limited by the pupil size. $Defo(u, v; z)$ is the angular spectrum defocus kernel and takes the form:

$$Defo(u, v; z) \;=\; e^{ikz\sqrt{1-(NA/n)^2}}, \tag{S2}$$

where *k* is the wavevector, $NA = \lambda\sqrt{u^2 + v^2}$ is the pupil coordinates relative to the numerical aperture, and *n* is the index of refraction of the propagation media. $A(u, v)$ is the on-axis Seidel aberrations characterized by our Zemax simulation (see **Supp. 1.2**).Since we are only considering third order on-axis aberrations, this term consists solely of spherical aberration . The trailing term is the expected intensity decay due to scattering.

Once we receive an optimized DOE, we may formulate a simulated image of a neuronal scene with a modified version of S1:

$$I_{im}(x, y) \;=\; \sum_z |F\{F\{o_{im}(x, y; z)\} \times M(u, v) \times D(u, v; z) \times A(u, v)\}|^2 \times e^{-z/l_s} + \eta(x, y), \tag{S3}$$

where $o_{im}(x, y; z)$ is now randomly distributed emitters on each slice. To simulate the strong out of focus background common in 1P neural imaging, $o_{im}(x, y; z)$ contains both 5µm target emitters and 1 µm background emitters that provide background fluorescence[2]. $\eta(x, y)$ is Gaussian noise applied to the resulting image that simulates the effects of sensor noise . When the optical signal exceeds 5 photons per pixel, shot noise can be well approximated as Gaussian noise and all common noise sources, excluding fixed pattern noise and flicker noise, can be combined into a single Gaussian noise source[3]. For the sake of simplicity, we assume that our sensor produces an ideal flat field so we neglect the effects of hot pixels and fixed pattern noise in simulation.

##### 1.2 System Characterization in Zemax

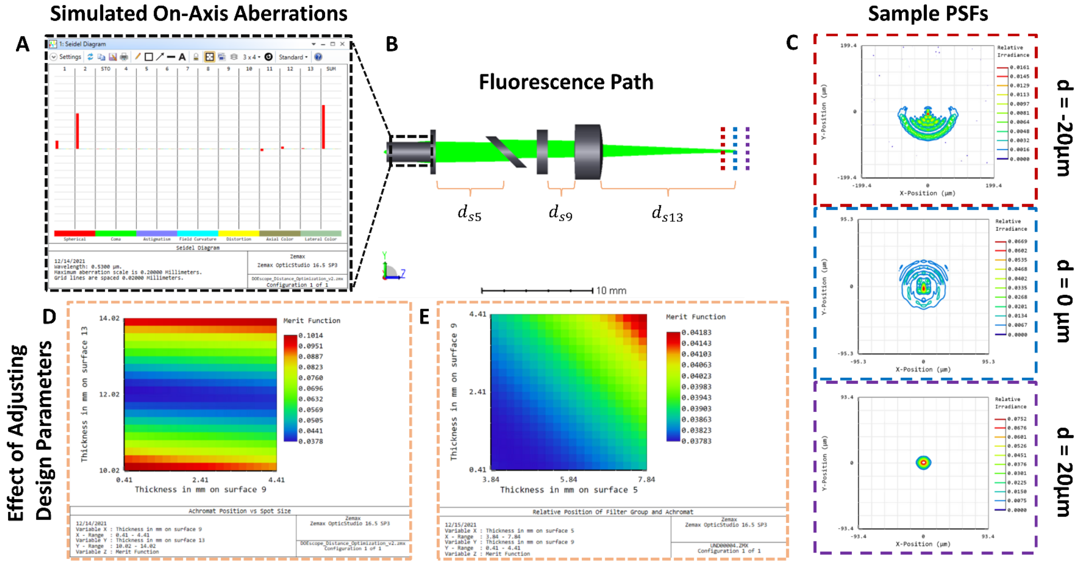


**Fig. S1.2. Zemax model of a EDOF-Miniscope with Proxy Phase Mask.** (**A**) On-axis seidel aberrations within an EDoF-Miniscope characterized by Zemax. (**B**) The shaded model of the emission/fluorescence path. The emitted optical signal enters the GRIN lens, passes through the glass substrate for the DOE, dichroic mirror and emission filter before being focused onto the imaging plane by the achromat. The simulation considers different distances between the optical components to explore their effect of the resulting spot size as a way of exploring the design space for the miniscope. (**C**) Sample PSFs predicted by the Zemax model at a few axial planes without the effects of the DOE. (**D**) 2D universal plot detailing the effect of changing the distance between the emission filter and achromat (surface 9) vs the achromat on the imaging plane (surface 13) on our merit function (rms spot size). (**E**) 2D universal plot detailing the effect of changing the distance between the GRIN lens and dichroic mirror (surface 5) vs the emission filter and achromat (surface 9) on our merit function (rms spot size).

Our Zemax model predicts the behavior and aberrations of the EDoF-Miniscope in free space using spectral information prepared by the relevant manufacturer (Chroma, Edmund Optics) in the sequential mode (see **Fig.S1.2A-B**). We exclusively examine the effects of the emission channel on the resulting PSF and assume that we have an ideal illumination channel (see **Fig.S1.2C**). We decouple these components so that we may explore the effects of the EDoF-Miniscope's design on the resulting PSF. The illumination channel will be dominated by the performance LED light source as collimated by an off-the-shelf ball lens. We first set the GRIN, achromatic lens and camera plane based on the effective focal lengths of each component as provided by the manufacturer. We next place an on-axis monochromatic source on the front focal plane of the GRIN lens with a central wavelength at the center of the GCamP spectrum (∼509nm) to optimize the distances between the emission channel optics (see **Fig.S1.2D-E**). We anticipate that any chromatic aberration will be negligible since our sources are bandpass filtered. We utilize a 2D universal plot to compare the effect of adjusting multiple components on the RMS spot size of our PSF in a compact plot. Adjusting the position of the emission filter had a negligible effect on the RMS PSF and adjusting the achromat could be compensated for by moving the imaging plane. As a result, future iterations of the EDOF-Miniscope may be shrunk without severely detrimenting the optical performance.

Importantly, Zemax predicts that, due to the strong spherical aberration, the PSF focuses before the nominal imaging plane. This reinforces our selection of a 0.23-mm working distance GRIN lens as opposed to a 0-mm working distance GRIN lens for fixed samples since the latter would truncate the focus on the platform. In our analysis, we replace the phase mask with a 0.5-mm thick glass cover slip to analyze the refractive effects of our DOE's substrate.

#### 2 Genetic Algorithm Convergence

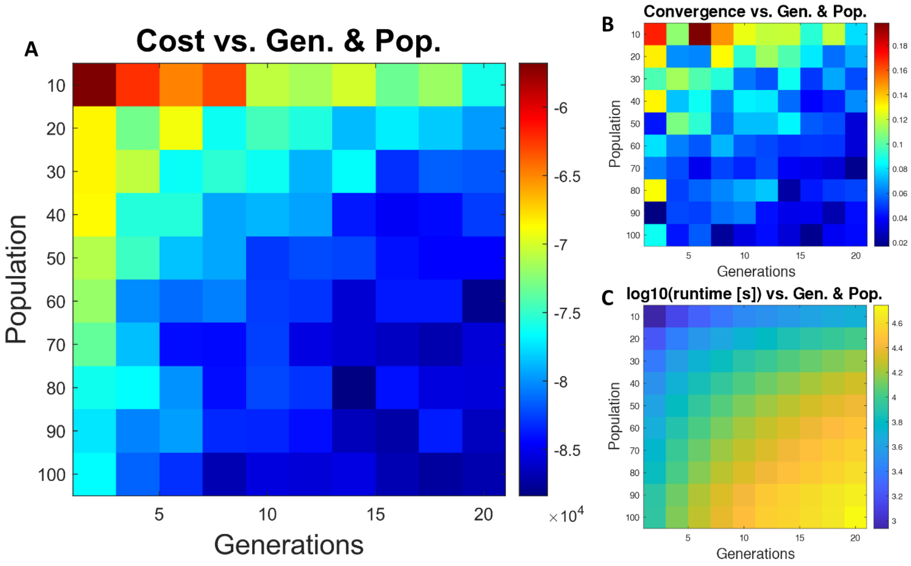


**Fig. S2.1. Convergence Analysis of the Genetic Algorithm.** (**A**). Analysis of the final cost of our optimized mask and undergoing optimization for a given population size for a certain number of generations. For robustness, we ran each trial 10 times and took the mean cost. (**B**) Here, we analyzed the coefficient of variation for each trial. In general, the coefficient decreases as population and generations increase, indicating that the genetic algorithm is achieving more stable solutions by increasing these parameters. (**C**) As population and generations increase, the optimization takes longer time to complete. To analyze the resources, we analyze the log of the runtime of the CPU to run a certain trial 1 time. While not shown, the algorithm achieves a comparable runtime over each of the 10 iterations each trial undergoes.

Genetic algorithms navigate complex landscapes, such as designing 3D sculpted light distributions[4] , by using the principles of evolution to iteratively refine the desired parameters within a population of candidate solutions using a user-crafted objective function. Often modeled as Markov Chains, prior work has shown that genetic algorithms are ergodic under weak conditions and will not remain in a suboptimal minima indefinitely regardless of the objective function or initial conditions. However, genetic algorithms often undergo punctuated equilibrium, which can make it difficult to quantize convergence[5] . We characterize the convergence properties of our genetic algorithm by sweeping through the population size and number of generations and analyzing the mean fitness value of the optimized map over 10 iterations (see **Fig. S2.1A**). We analyze the coefficient of variation and average optimization time to better interrogate the tradespace between convergence and computational cost (see **Fig. S2.1B-C**).

We reasonably converge near a stable optimum in a small number of iterations (~10) with a moderate sized population of 60 candidate DOEs. While this optimization does inflate computation time, our genetic algorithm is still able to optimize a DOE in 80 minutes. This analysis reinforces our choice in using a genetic algorithm as a fast paced solution for optimizing an EDoF-Miniscope for deployment in scattering media.

### 3 Experimental Test Setups

#### 3.1 High Precision DOE Alignment Setup

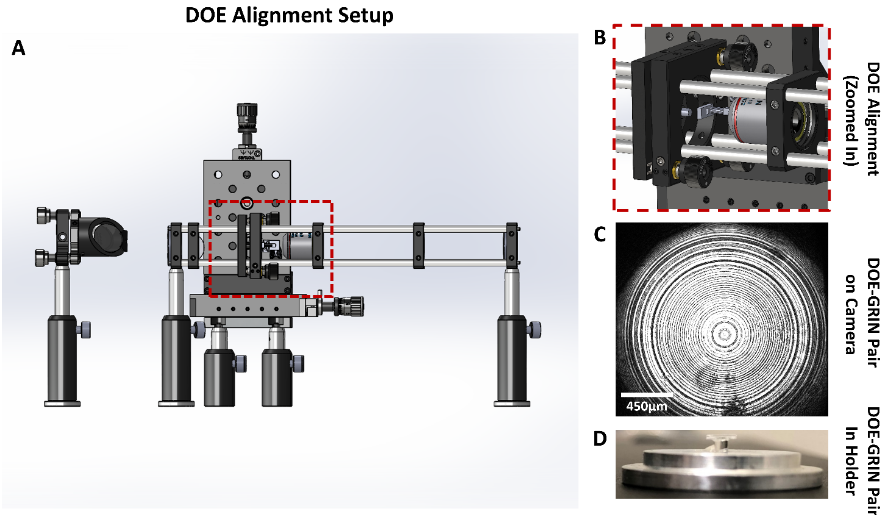


**Fig. S3.1. Precision Alignment of the DOE and GRIN Lens.** (**A**) Optical setup used to align the DOE and GRIN lens. The cage mounted setup aligns the laser illumination source to the GRIN lens. The free space stage brings the DOE onto the optical axis and into contact with the GRIN lens to adhere them together. (**B**) Zoom in on the binary DOE holder and GRIN lens to showcase the alignment procedure. (**C**) A final aligned DOE-GRIN pair as viewed from the visual feedback camera. All alignment is done under visual guidance by adjusting the DOE relative to the fixed GRIN lens. (**D**) A final aligned DOE-GRIN pair after removing it from the alignment setup.

We align the DOE to the GRIN lens using the setup shown in **Fig. S3.1A-B**. First, a collimated laser, after passing through a spatial filter (Newport, 910A) with a 10x objective (Newport, M-10X) before passing through a 25 micron pinhole (Newport, 910PH-25) and is then deflected by an elliptical mirror (Thorlabs, BBE1-E02) in a 45 degree mount (Thorlabs, H45E1) into the cage mounted setup. The cage mount contains two 4f optics systems: the miniaturized optics system and a relay optics system. The miniaturized optics system consists of a primary lens (Thorlabs, f= 25mm, LA1951), the GRIN lens mounted in a 1 inch optics holder (Thorlabs, CP33) in a custom milled mount (see **Fig. S3.1D**) and the DOE. The primary lens and GRIN lens act as a 4f system to demagnify the laser and illuminate the DOE, which is integrated into the setup through a custom holder (See **Fig. S3.1B**) and a 3-axis linear stage (Thorlabs, PT3A) for high precision positioning. The relay system consists of an objective (Nikon, MRL00042) and tube lens (Thorlab, f = 150mm, LA1433) and magnifies the resulting image onto a camera

(FLIR, BFLY-PGE-50A2M-CS) (see **Fig. S3.1C**) to enable visual feedback for high precision alignment.

To adhere the DOE to the GRIN lens, we use the following procedure:

1. Remove all components to the left of the objective lens as shown in Fig. 3A except for the elliptical mirror and mount.
2. Remove the GRIN holder from the CP33 O1” optics mount.
3. Place a mask blank in the back slot of the GRIN holder.
4. Place the GRIN in the through hole and adhere it in place using a small droplet of dental paste.
5. Determine the top surface of the DOE (i.e. the face with the features etched in it). We use KLA Tencor Alpha Step 500 Surface Profiler, which has an oblique illumination channel and allows us to visually determine the top from bottom surface of a finished DOE.
6. Place the DOE in the DOE holder so that the surface with the pattern faces away from the GRIN lens.
7. Attach the DOE holder onto the PT3A so that it is roughly on-axis and in front of the objective lens.
8. Turn on the laser illumination and use the PT3A to bring the mask in-focus in front of the objective using visual guidance.
9. Scan the DOE along z, away from the objective lens, to ensure all components are well aligned on the optical axis and there is no noticeable drift in the DOE position on the camera.
10. Place the GRIN holder back in the CP33 mount and rebuild all components to the left of the GRIN lens. Use a level to ensure the GRIN lens is on the optical axis.
11. Physically place the CP33 mount close to the DOE such that the DOE is within the aperture of the GRIN lens but not so that the components are touching.
12. Place a small droplet of NOA63 glue on the exposed surface of the GRIN lens.
13. Use the PT3A stage to bring the DOE in contact with the GRIN lens. Since the GRIN lens might now be perfectly straight, use the x- and y-axes to keep the DOE in the center of the GRIN’s aperture.
14. After achieving contact, make any last adjustments and cure the glue by illuminating the DOE-GRIN pair through the objective with the UV source.
15. Remove the CP33 Mount with the now adhered DOE - GRIN pair and integrate into an EDOF-Miniscope.

##### 3.2 Automated Test Platform for Fixed Samples

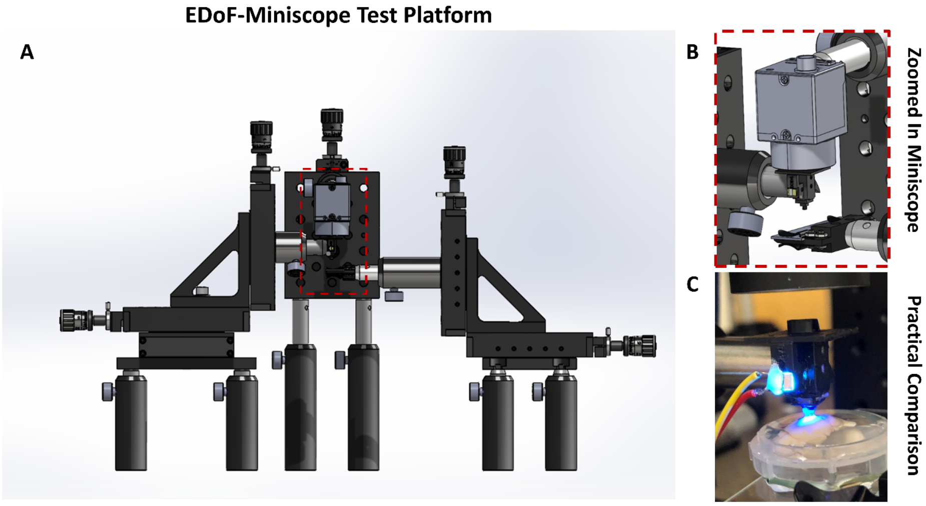


**Fig. S3.2. Automated Test Platform for Static Samples.** (**A**). The test setup consists of two three-axis thorlabs linear 1" stages (PT3A, Thorlabs) to hold the miniscope and sample respectively as well as one one-axis linear 1" stage (PT1A, Thorlabs) for the camera. The z-stage of the sample stage has been automated through Pycromanager to synchronize stage movement with camera acquisitions. (**B**) To achieve a high precision alignment, the EDOF-Miniscope is mounted on an O1/2" thorlabs post and manually aligned to the sample using the camera as a reference. (**C**) Example of the alignment of the EDOF-Miniscope to the whole fixed mouse brain as a practical example.

We integrated the EDoF-Miniscope into the setup described in **Fig. S3.2A** to perform our experiments on fixed samples. The setup consists of two linear 3-axis stages (Thorlabs, PT3A), and a linear 1-axis stage (Thorlabs, PT1A). One three axis stage adjusted the xyz position of the EDoF-Miniscope through a thorlabs optical post fixed into the side fin of the platform. The other three axis stage was outfitted with a sample holder (Thorlabs, MAX3SLH) to adjust the XYZ position of several fixed samples on glass slides. The one-axis stage adjusted the z position of the camera. To align the camera and miniscope, we first placed a sample of sparse 1µm fluorescent beads under the EDoF-Miniscope and adjusted the position of the miniscope and camera until the beads were in focus, as depicted in **Fig. S3.2B.** Next, we replaced the 1µm fluorescent beads sample with another fixed sample and used the sample stage to bring the new sample into focus. The z-axis of the sample stage was replaced with a DC actuator (Thorlabs, Z806) and automated through Pycro-Manager[6] . This allowed us to automate both the position of our sample as well as our camera acquisition to synchronize our data collection. We utilized this platform to capture single plane as well as z-scan imaging sessions of fixed samples.

#### 4 DOE Perturbation Analysis

##### 4.1 Effect of Lateral and Axial DOE Displacement on the DoF

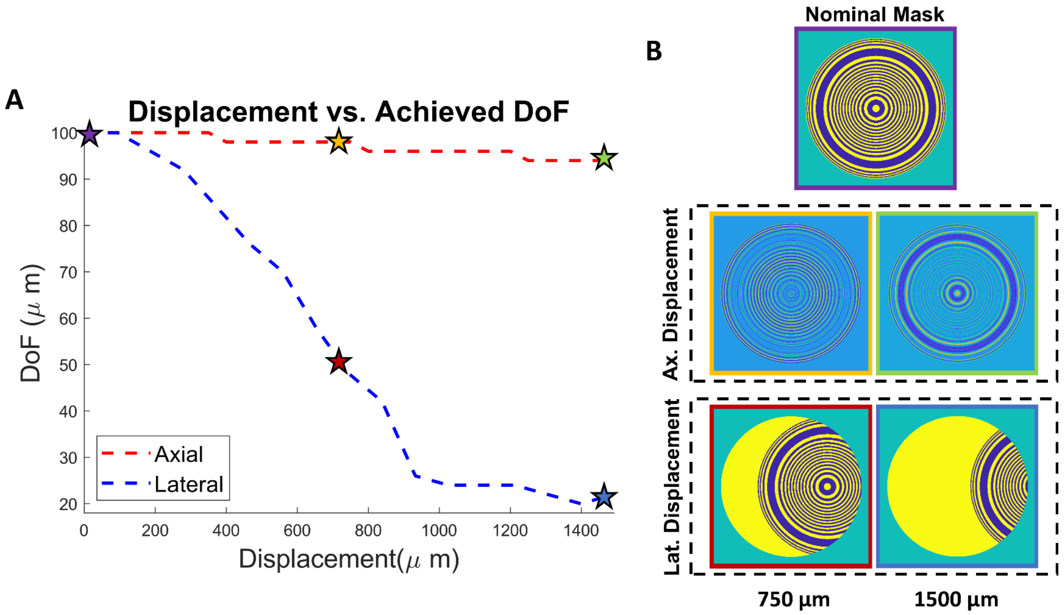


**Fig. S4.1. Effect of Lateral and Axial DOE Displacement on the DoF.** (**A**) Achievable DoF by our EDoF-Miniscope when the DOE is displaced axially (red line) and laterally (blue line) on the pupil plane. (**B**) Examples of the resulting pupil function without modification (top row), when the DOE is axially displaced by 750 µm and 1500 µm (middle row) and when laterally displaced by 750 µm and 1500 µm (bottom row).

We characterize the effects of axially and laterally displacing our DOE on the pupil plane on our DoF in simulation to analyze our alignment tolerances when assembling the EDoF-Miniscope in **Fig.S4.1A**. We utilize (S1) to simulate the 3D intensity distribution, where *M(u,v)* is replaced by our displaced mask. To perform lateral displacement, we first utilize the Matlab circshift command in Matlab to shift our mask on our pixelated grid by the nearest integer number of pixels to our desired displacement amount. To perform axial displacement, we utilize the defocus kernel described in (S2) to simulate the optical field generated by an axially displaced mask when back-propagated to the pupil plane. We find that axial displacement is a weak parameter on the resulting DoF. We predict that this is due to the fact that, in a miniscope, any axial displacement will be small compared to the size of the DOE. In addition the rings that comprise the DOE vary in size between ~4 µm and ~150 µm. While small rings may blur as the DOE defocuses, **Fig.S4.1B (middle)** shows that many of the larger structures are still perceivable within our simulated range.

We find that the mask is more sensitive to lateral displacement. Intuitively, we may model small lateral shifts as convolutions of our DOE with an off-axis delta function, which will result in sheared versions of our PSF in any cross-sectional plane but should not significantly affect the achieved DoF. However, as we continue to laterally move the DOE, the DOE pattern is increasingly clipped by the aperture and replaced by the nominal bare aperture of the GRIN lens (**Fig.S4.1B (bottom)**). As this occurs we expect the DoF to significantly decrease and ultimately converge to the nominal DoF, which is 20 µm for this given simulation. From this analysis we may conclude that our DOE is negligibly affected by our reported lateral alignment accuracy (70 μm) and any deviation between the thickness of our substrate (500 μm) and working distance (230 μm).

We simulated all displacements for fixed parameters: $n$ = 1.33, $l_s$ = 100 μm, for 5 μm on-axis sources, over a 120 μm range with a 2 μm step size. We characterize DoF using the number of depth planes that contain an intensity value within 50% the maximum 3D value.

##### 4.2 Alignment Precision under Visual Guidance

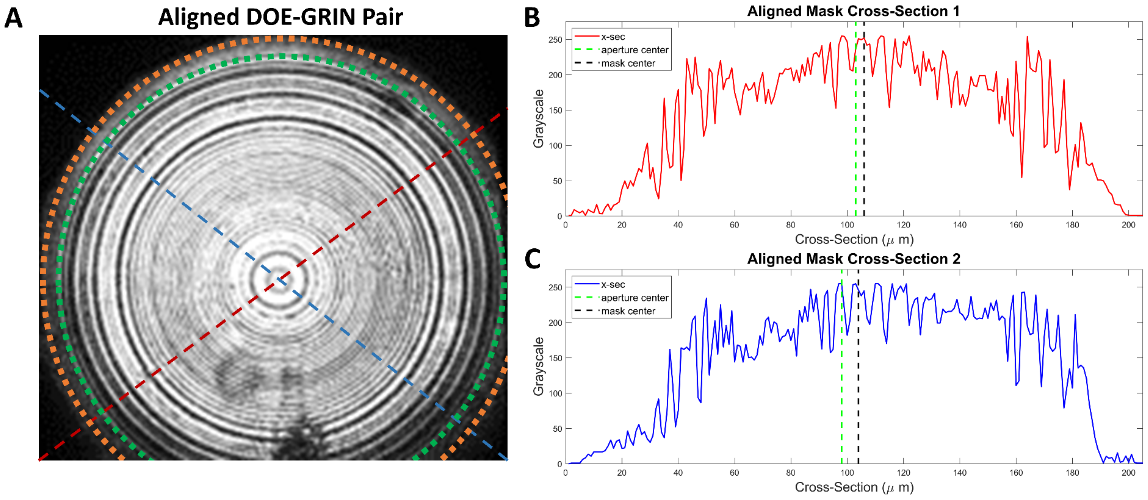


**Fig. S4.2. Achieved Precision under Visual Alignment.** (**A**). Example feedback image when visually aligning the DOE to the GRIN lens using the setup in Fig. 3A. The two cut lines correspond to (**B**) and (**C**) respectively. The GRIN circle marks the cutoff of the mask and the orange circle marks the cutoff of the aperture.

After performing visual alignment, we characterize the lateral precision of our alignment of our DOE - GRIN pair to confirm that our setup can achieve a precision within the tolerances described by **Supp. 4.2**. We characterize our alignment by comparing the geometric center of the DOE to the geometric center of the surrounding GRIN aperture (See **Fig. S4.2A**). The center of the DOE is easily identifiable by the structure of the center ring. We determine the cutoff of the aperture as the closest pixels on either side of the DOE that reach a grayscale value of 0. We can then determine the center of the aperture as the midpoint between those coordinates and compare that calculated value to the midpoint of the DOE to determine our alignment accuracy. For robustness, we perform that analysis over multiple cutlines (See **Fig. S4.2B-C**) to determine that we have a *maximum* lateral misalignment of 70μm. The material list for the high-precision alignment setup as well as CAD models for the custom components can be found on our GitHub.

#### 5 EDoF-Miniscope Characterization on Fluorescent Samples

##### 5.1 Axial Elongation Within a  5μm Scattering Phantom

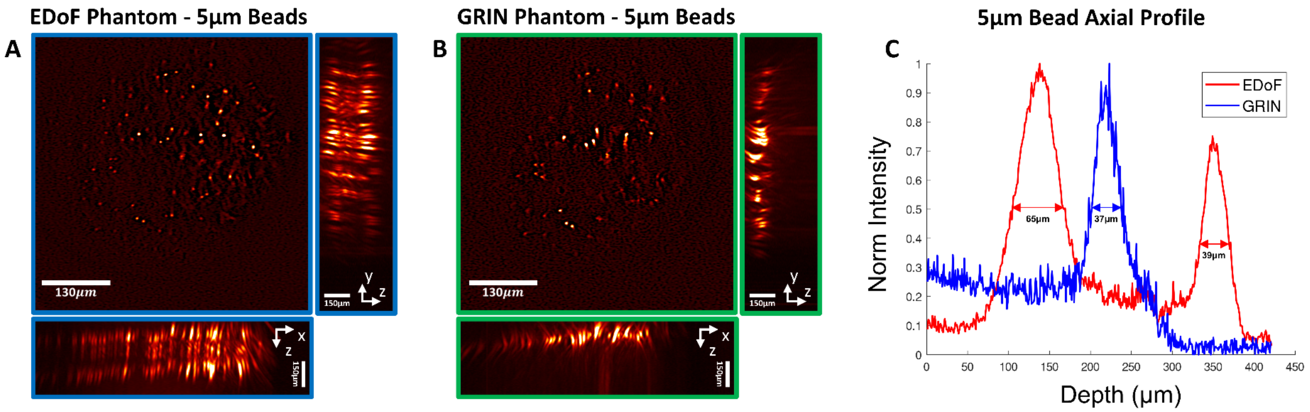


**Fig. S5.1. Axial Elongation Within Scattering Phantom with 5-µm beads.** (**A**) Volumetric information from an axial scan of a scattering phantom ($l_s$ = 100 µm) embedded with 5 µm beads imaged by the EDoF-Miniscope. The center image is one in-focus plane from the 1st diffraction order and the side and bottom images are xz and yz MIPs respectively. (**B**) Volumetric information from an axial scan of the same scattering phantom imaged by a miniscope. The center image is one in-focus plane from the 1st diffraction order and the side and bottom images are xz and yz MIPs respectively. (**C**) Axial profile from cut line on MIP for the EDoF-Miniscope and miniscope.

We characterize the axial elongation of 5 µm fluorescent beads in a scattering phantom ($l_s$ = 100 µm) as a proxy for neurons in neural tissue. After using the setup described in **Supp. 3.2** to bring the sample into focus for the EDoF-Miniscope and miniscope, we utilize an automated sample stage to acquire a z-stack. We show the one focal plane image combined with xz and yz MIPs for the EDoF-Miniscope (see **Fig. S5.1A**) and the miniscope (see **Fig. S5.1B**) respectively. We plot the cut lines from **Fig. S5.1A-B** in **Fig. S5.1C** to characterize the axial elongation. The EDoF-Miniscope achieves an axial elongation of 107 µm between the two foci while the miniscope achieves an axial elongation of 37 µm.

#### 5.2 Axial Elongation Within a Scattering Phantom with 10-µm beads

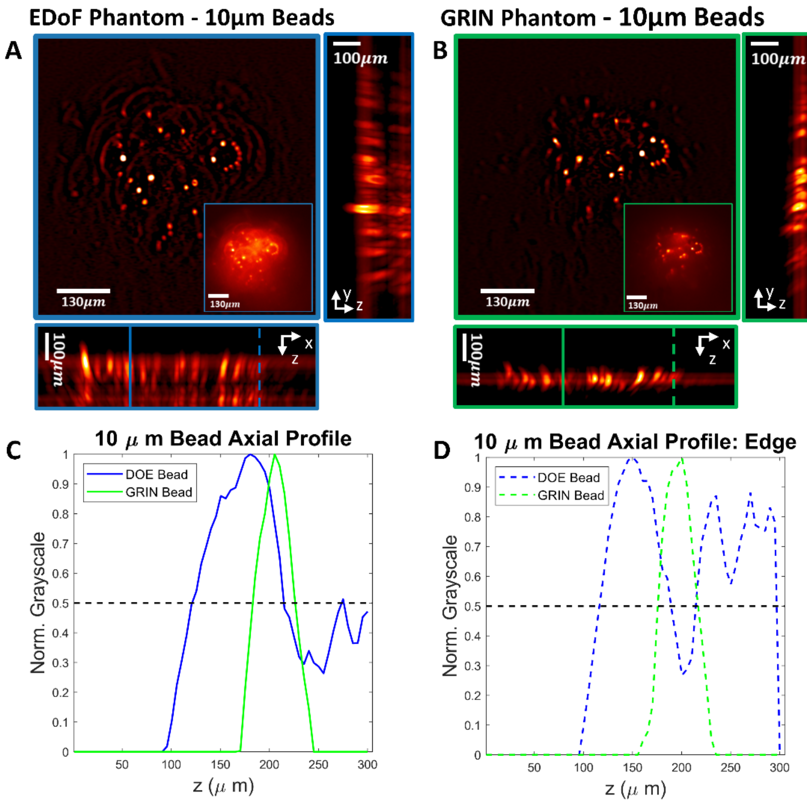


**Fig. S5.2. Axial Elongation Within a Scattering Phantom with 10-μm beads.** (**A**) Volumetric information from an axial scan of a scattering phantom ($l_s$ = 100 μm) with 10 μm beads imaged by the EDoF-Miniscope. The center image is one in-focus plane from the 1st diffraction order and the side and bottom images are xz and yz MIPs respectively. (**B**) Volumetric information from an axial scan of the same scattering phantom imaged by a miniscope. The center image is one in-focus plane from the 1st diffraction order and the side and bottom images are xz and yz MIPs respectively. (**C**) Cut lines near the center of the FoV from the EDoF-Miniscope and miniscope. (**D**) Cut lines near the edge of the FoV from the EDoF-Miniscope and miniscope.

We characterize the axial elongation of 10 μm fluorescent beads in a scattering phantom ($l_s$ = 100 μm) as a proxy for large neurons in neural tissue. After using the setup described in **Supp. 3.2** to bring the sample into focus for the EDoF-Miniscope and miniscope, we utilize an automated sample stage to acquire a z-stack. We show the one focal plane image combined with xz and yz MIPs for the EDoF-Miniscope (see **Fig. S5.1A**) and the miniscope (see **Fig. S5.1B**) respectively. We plot the cut lines from **Fig. S5.1A-B** in **Fig. S5.1C** to characterize the axial elongation. The EDoF-Miniscope achieves an axial elongation of 95 μm in the first order near the center of the FoV and 85 μm near the edge of the field of view. The miniscope achieves an axial elongation of 45 μm in the first order near the center of the FoV and 40 μm near the edge of the field of view

##### 5.3 SBR Analysis Over 5-μm Bead Phantoms of Varying Density

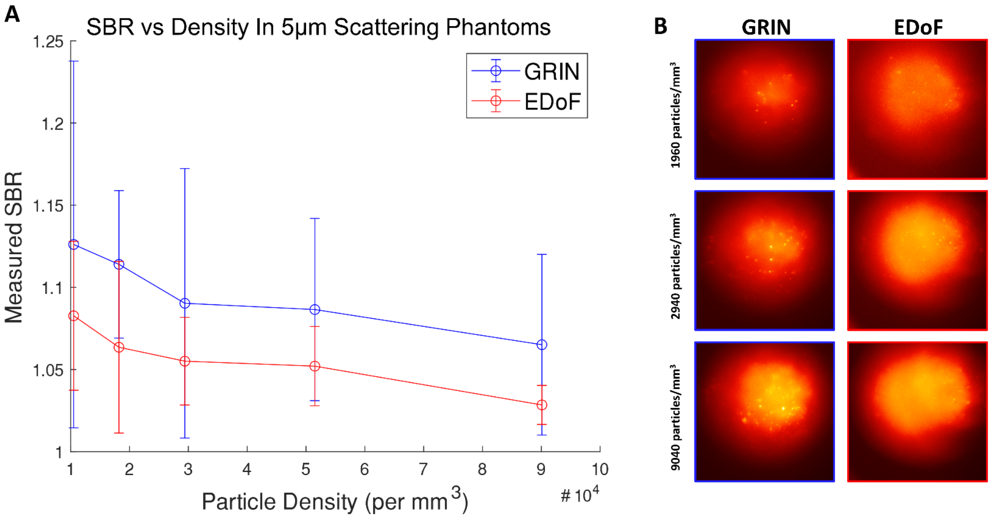


**Fig. S5.3. SBR Analysis Over 5μm Bead Phantoms of Varying Density.** (**A**) Signal-to-Background ratio averaged over 10 particles across the FoV of several phantoms of varying density as imaged by the EDoF-Miniscope and a miniscope. (**B**) Example images from the miniscope (left) and EDoF-Miniscope (right) of several phantoms of different densities.

To investigate any reduced contrast in the EDoF-Miniscope, we imaged 5μm bead scattering phantoms of controlled densities (~1000-10000 particles/mm$^3$) with both the EDoF-Miniscope and a miniscope. We randomly determined the centroids of 10 particles across the FoV of each image and determined the signal by averaging a 8 pixels circular region-of-interest around each centroid and determined the background by averaging a 35 pixels circular region-of-interest. The mean SBR and standard deviation were determined for each density (see **Fig. S5.3A**). In general, the EDoF-Miniscope exhibits a 4% decrease in SBR over all phantoms (see **Fig. S5.3B**).

### 6 Analysis of Wavelet Filter

#### 6.1 Simulated Wavelet Performance

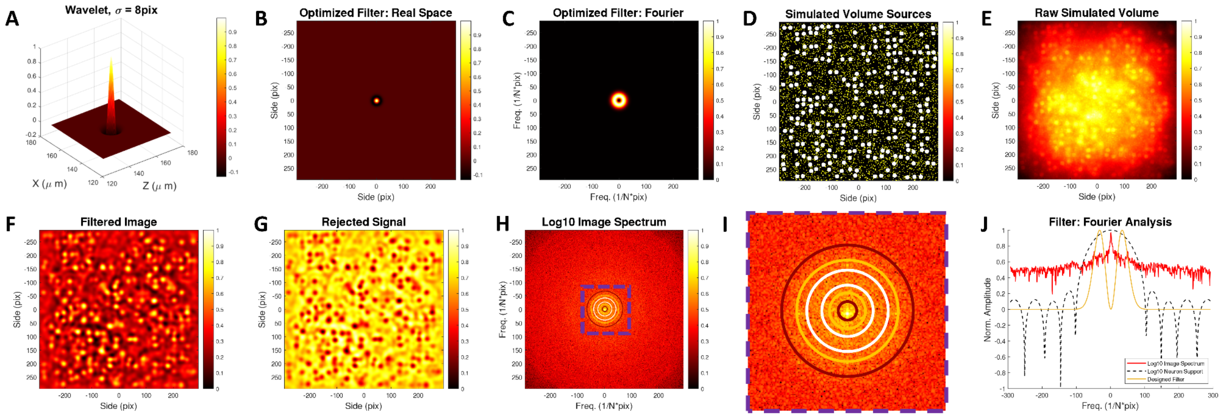


**Fig. S6.1. Simulated Wavelet Filter.** (**A**) A surface profile of the wavelet kernel utilizes in processing static neuronal frames. (**B**) Real space wavelet convolution kernel and (**C**) its profile in the fourier domain. (**D**) The MIP of a simulated test volume consisting of 5 μm neurons and 1 μm background particles. (**E**)

The simulated raw measurement when the simulated volume was distributed over 150 µm. (**F**) The wavelet process framed and (**G**) the rejected signals from the wavelet transform. (**H**) An overlay of the wavelet transform Fourier spectrum (contour) on the spectrum of the simulated image (image) with (**I**) a zoom-in on the spectrum and (**J**) a cross-section of the image spectrum, Fourier space wavelet and the spectrum of a 5-µm proxy particle.

We analyze the performance of the wavelet used for post-processing our fixed samples, following a proposed form[7] (see **Fig. S6.1A**). We found that utilizing a Laplacian of Guassian (LoG) wavelet with a standard deviation of 8 pixels performs adequate background suppression while extracting neuronal structures (see **Fig. S6.1B**). Intuitively, the LoG wavelet is an annulus in Fouirer space, denoting its ability to suppress constant and slowly varying background as well as select objects within a certain band (see **Fig. S6.1C**). We simulated a neuronal volume while imaging with an EDoF-Miniscope with 5 µm proxy neurons with 1 µm sources to artificially add background (see **Fig. S6.1D-E**). Applying the wavelet transform well extracts the 5 µm proxy particles while suppressing the background and 1 µm sources (see **Fig. S6.1F-G**). We overlay the Fourier transform of the designed wavelet filter on the spectrum of the simulated image to emphasize that the wavelet is tailored to extract the target particles (see **Fig. S6.1H-J**).

##### 6.2 Comparison to Deconvolution - Brain Slice Sample

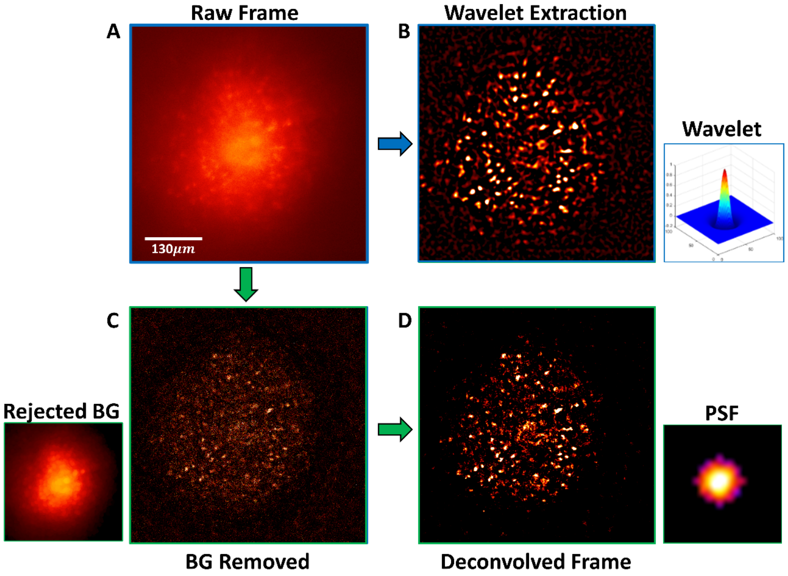


**Fig. S6.2. Comparison of Wavelet Filtering to Deconvolution on Neural Slices.** (**A**) Raw image acquired by imaging a 100 µm fixed mouse brain slice with an EDoF-Miniscope. (**B**) EDoF-Miniscope image after being processed by wavelet filtering. (**C**) EDoF-Miniscope image after morphological background removal and (**D**) deconvolution.

We show that wavelet filtering visually outperforms standard deconvolution on a 100 µm fixed mouse brain slice. We first capture a raw frame of a 100 µm fixed mouse brain slice by the EDoF-Miniscope (see **Fig. S6.2A**). We process the raw frame with a LoG wavelet with a standard deviation of 8 pixels (see **Fig. S6.2B**). We compared the wavelet filtered image to an image that was post-processed with morphological background removal[8] and deconvolution with the PSF from **Fig.1D** and Tikhonov regularization (see **Fig. S6.2C-D**). Wavelet filtering performs more effective extraction of the fixed neurons across a larger FoV when compared to the deconvolved image. We predict that the wavelet filter is effective in two parts. First, the wavelet filter is summable to zero, allowing it to more effectively reject high background. Second, the wavelet filter exhibits a gaussian peak which averages several pixels and reduces noise. Altogether, these properties allow the wavelet filter to effectively extract neuronal signals from low contrast, high noise images.

##### 6.3 Comparison to Deconvolution - Vasculature

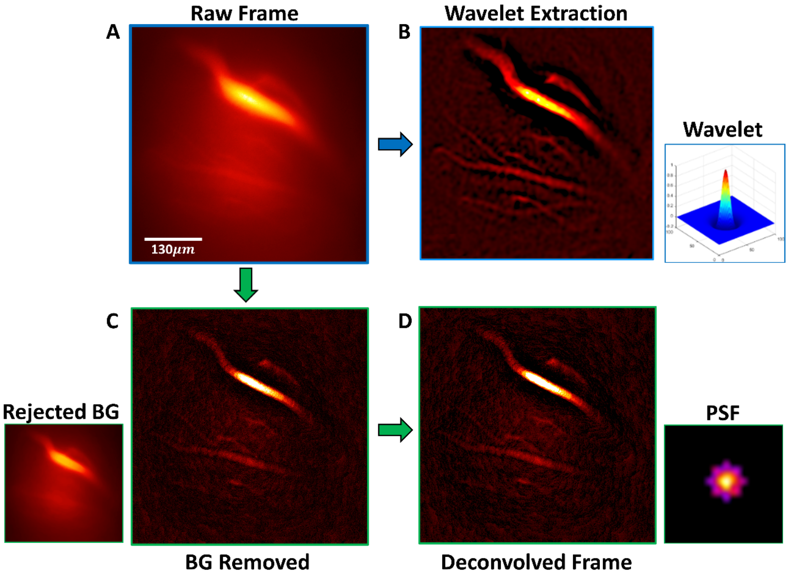


**Fig. S6.3. Comparison of Wavelet Filtering to Deconvolution on Stained Vasculature.** (**A**) Raw image acquired by imaging stained vasculature in a whole mouse brain with an EDoF-Miniscope. (**B**) EDoF-Miniscope image after being processed by wavelet filtering. (**C**) EDoF-Miniscope image after morphological background removal and (**D**) deconvolution.

We show that wavelet filtering visually outperforms standard deconvolution on stained vasculature in a whole fixed mouse brain. We first capture a raw frame with the EDoF-Miniscope (see **Fig. S6.3A**). We process the raw frame with a LoG wavelet with a standard deviation of 8 pixels (see **Fig. S6.3B**). We compared the wavelet filtered image to an image that was post-processed with morphological background removal and deconvolution with the PSF from **Fig.1D** and Tikhonov regularization (see **Fig. S6.3C-D**). Wavelet filtering performs more effective extraction of the vasculature across a larger FoV when compared to the deconvolved image. Notably, the morphological background removal and deconvolution fail to recover the entirety of the small vasculature present in the bottom of the image and artificially constrain the width of the large vessels at the bottom of the image. This trial emphasizes that wavelet filtering is an effective tool at extracting a variety of neural signals and generalizes across several source geometries.

#### 7 Exploration of DOE Optimization In Different Scattering Media

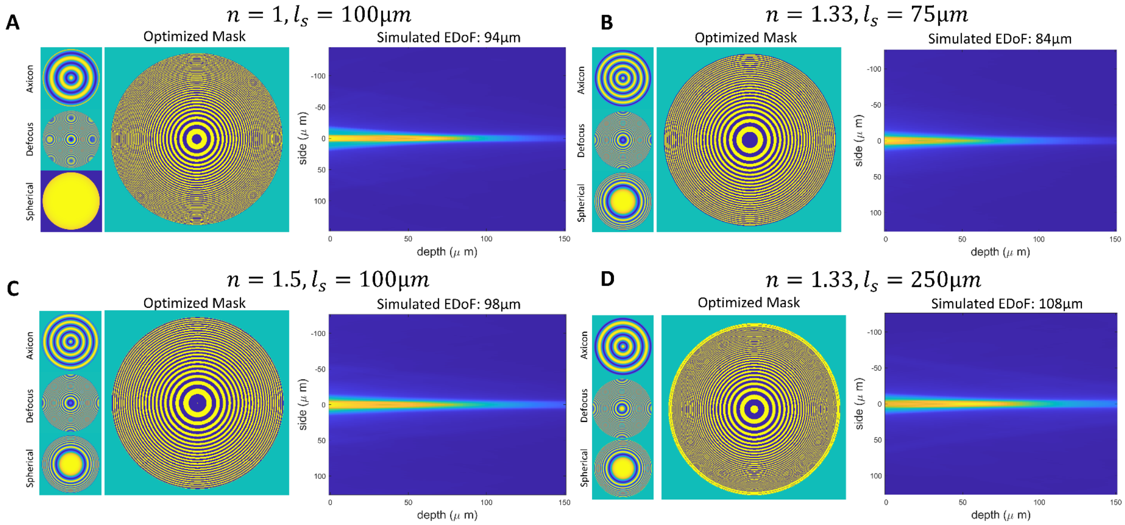


**Fig. S7.1. DOE Design for Different Scattering Media.** The learned optimized phase basis (left), binarized mask (center) and the 1st diffraction order elongation (right) for a media with **(A)** an index near air ($n$ = 1) and similar scattering to our original case ($l_s$ = 100 µm). (**B**) an index near water ($n$ = 1.33) and stronger scattering to our original case ($l_s$ = 75 µm). (**C**) an index near oil ($n$ = 1.5) and similar scattering to our original case ($l_s$ = 100 µm). (**D**) an index near water ($n$ = 1.33) and similar scattering to our original case ($l_s$ = 250 µm).

To explore the generalizability of our genetic algorithm, we explore the optimization of several DOEs under different designed conditions. Since our genetic algorithm directly tailors the first diffraction order and our future work will utilize zero working distance GRIN lenses, we judge the achieved EDoF in the first order after optimization. All optimizations assumed that we are elongating a 5 µm on-axis source and aimed to achieve an elongation of 100 µm. We first analyze the effect of changing the assumed index of refraction while retaining a comparable scattering length to our original design (See. **Fig. S7.1A,C**). In both cases, we are able to achieve elongations near the target depth. Next we fixed the index and changed the scattering length to shorter than (See **Fig. S7.1B**) and longer than (See **Fig. S7.1D**) the original design. While, as expected, the DOE designed for a longer scattering length achieves the desired axial elongation the DOE designed for a shorter scattering length was unable to achieve an elongation significantly past one scattering length. Intuitively, this makes sense as the optical intensity decays to under 36% after one scattering length, increasing the difficulty to maintain a relative intensity within 50% of the peak value (our definition of axial elongation). As a result, we confirm that our genetic algorithm is well generalizable for different media however is fundamentally limited by the scattering length. It should be noted that the mask optimized in air unlearned spherical aberration while all other masks learned a unique combination of all three basis functions. This indicates that the optimal mask is depending on the user defined conditions and no one mask generalizes across different cases. This reinforces the need for a rapid algorithm that can optimize DOEs for EDoF imaging across a variety of applications.
